# Comprehensive characterisation of molecular host-pathogen interactions in influenza A virus-infected human macrophages

**DOI:** 10.1101/670919

**Authors:** Sara Clohisey, Nicholas Parkinson, Bo Wang, Nicolas Bertin, Helen Wise, Andru Tomoiu, FANTOM5 Consortium, Kim M. Summers, Piero Carninci, Alistair A. Forrest, Yoshihide Hayashizaki, Paul Digard, David A. Hume, J. Kenneth Baillie

## Abstract

Macrophages in the lung detect and respond to influenza A virus (IAV), determining the nature of the immune response. Using terminal depth 5’-RNA sequencing (CAGE) we quantify transcriptional activity of both host and pathogen over a 24-hour timecourse of IAV infection in primary human monocyte-derived macrophages (MDM). We use a systems approach to describe the transcriptional landscape of the host response to IAV contrasted with bacterial lipopolysaccharide treated MDMs, observing a failure of IAV-treated MDMs to induce feedback inhibitors of inflammation. Systematic comparison of host RNA sequences incorporated into viral mRNA (“snatched”) against a complete survey of background RNA in the host cell enables an unbiased quantification of over-represented features of snatched host RNAs. We detect preferential snatching of RNAs associated with snRNA transcription and demonstrate that cap-snatching avoids transcripts encoding host ribosomal proteins, which are required by IAV for replication.

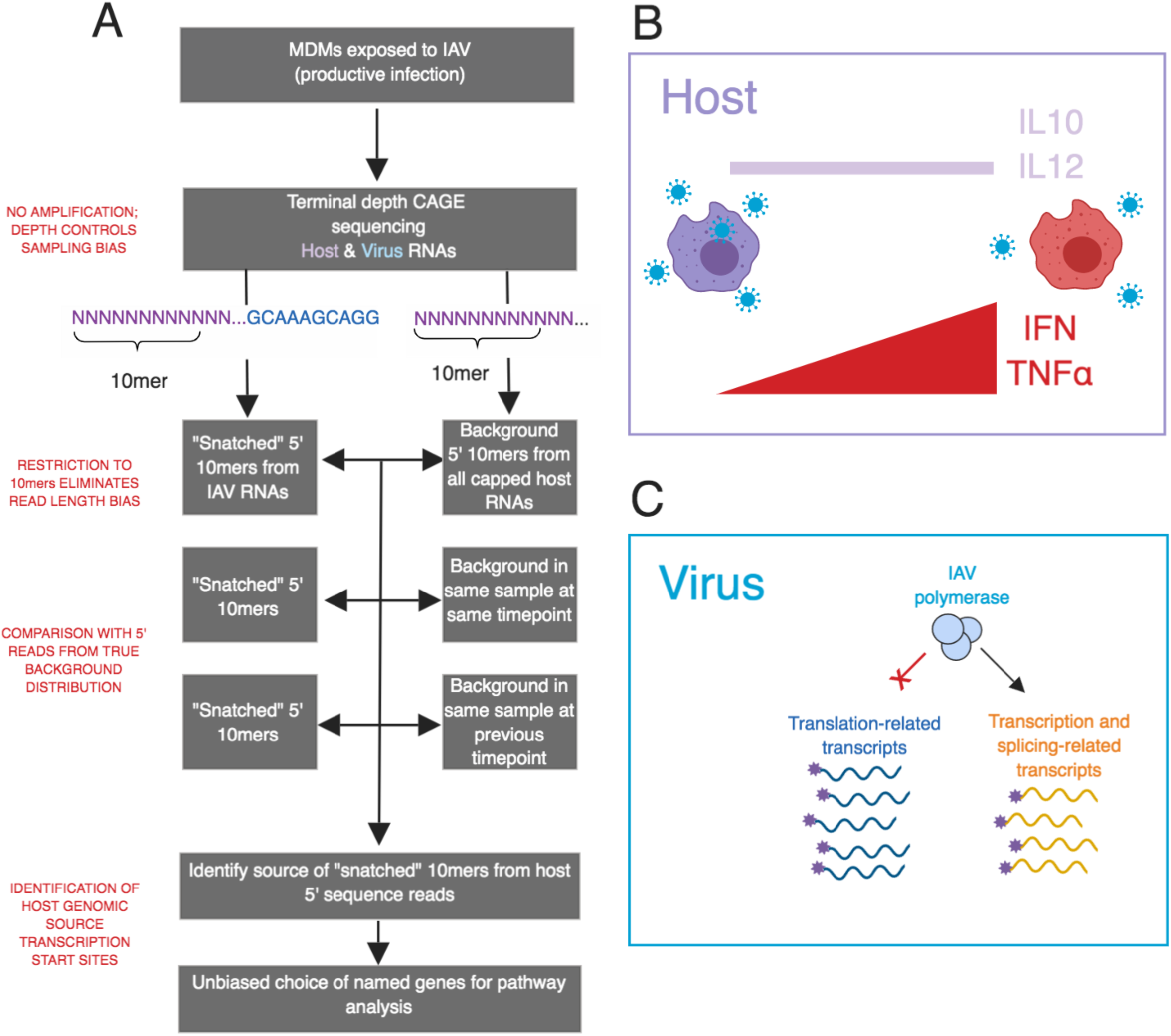

**Graphical Abstract:** (A) Overview of bioinformatics pipeline. (B) Host gene expression reveals that human macrophages exposed to IAV exhibit sustained production of key inflammatory mediators and failure to induce expression of feedback inhibitors of inflammation. (C) Unbiased comparison with total background RNA expression demonstrates that IAV cap-snatching has a strong preference for, and aversion to, different groups of host transcripts.

## Introduction

Influenza A virus (IAV) infection is responsible for an estimated 500,000 deaths and up to 5 million severe respiratory illness cases each year (WHO). The virus infects the respiratory tract, binding to and infiltrating the respiratory epithelium. The abundant macrophages of the airway and lung interstitium detect and respond to the virus, determining both the nature and the magnitude of the innate and acquired immune response (Cline, Beck and Bianchini, 2017) and contributing to inflammatory cytokine production outside the lung in severe IAV (Short *et al*., 2017). Human monocyte-derived macrophages (MDM) can be infected with IAV, produce viral proteins and release inflammatory cytokines in response to infection, thus they have been widely-studied as an experimental model (Hoeve *et al*. 2012; Perrone *et al*. 2008; van Riel *et al*. 2011; Monteerarat *et al*. 2010; Stasakova *et al*. 2005; Wang *et al*. 2012; Lee *et al*. 2009). In most studies, infected macrophages have produced relatively small yields of infectious IAV, although this has differed depending upon the virus strain and its virulence, and the cell population studied (Perrone *et al*. 2008; Stasakova *et al*. 2005; Wang *et al*. 2012; Friesenhagen *et al*. 2012; Nicol and Dutia 2014).

IAV is an RNA virus, containing 8 negative-sense segments that are transcribed and replicated in the nucleus of the host cell. As an obligate intracellular parasite, IAV is reliant on host cellular machinery for replication. To accomplish mRNA production, the IAV polymerase binds directly to the 5’ 7-methylguanylate cap of a nascent host RNA and cleaves it roughly 10-14 nucleotides downstream. The snatched sequence, known as a “leader” sequence, is employed as a primer for efficient transcription of the viral mRNA (Plotch *et al*., 1981) and provides the cap to viral mRNA to facilitate translation by host ribosomes. Previous large-scale studies of this process (Gu *et al*., 2015; Koppstein *et al*., 2015; Sikora *et al*., 2017, 2014) have produced evidence that host-derived RNA caps are frequently snatched from non-coding RNAs, particularly small nuclear RNAs (snRNAs), due to their high abundance in infected cells. This has led to the conclusion that cap-snatching is not a selective process – that is, that host mRNAs are snatched at random (Sikora *et al*., 2017; De Vlugt, Sikora and Pelchat, 2018). These previous RNA-Seq studies have detected snatched leaders, but have been unable observe the complete pool of unsnatched sequences, because of limited sequencing depth and resolution at the 5’ end, both of which are necessary to accurately quantify the background distribution of each host transcript. The CAGE RNA sequencing method captures both host and virus-derived transcripts and, importantly, does not require a PCR amplification step, thus eliminating PCR bias.

To overcome these limitations, we utilised cap analysis of gene expression (CAGE) to sequence capped RNA from the primary MDMs of 4 human donors *in vitro* at 4 time points over the course of a 24 hour, productive infection with IAV. This allows us to observe the transcriptional response to IAV infection over time in unprecedented molecular detail. This work was carried out as part of the FANTOM5 consortium. Data are accessible through the FANTOM5 ZENBU browser (http://fantom.gsc.riken.jp/zenbu/) and the FANTOM5 Table Extraction Tool (http://fantom.gsc.riken.jp/5/tet/).

We employ a systems approach to identify key features of transcription during IAV infection in MDMs. We previously used CAGE to quantify, transcript expression, promoter and enhancer activity in human MDM and produced a detailed time course profiling their response to bacterial lipopolysaccharide (LPS) (Baillie *et al*., 2017). As in our previous work, we use the principle of coexpression to identify key biological processes (Forrest *et al*., 2014; Baillie *et al*., 2016), and compare the response of MDMs to both IAV and LPS, revealing IAV-specific features of the host response.

By comparing the sequences of the snatched population to the sequences of the total capped RNA background, we have identified nucleotide sequence motifs associated with viral cap snatching and, for the first time, motifs present in the background host mRNA population that are not snatched. Furthermore, we have assigned transcript identity to leader sequences and observed potential preferential snatching of transcripts encoding spliceosome components and avoidance of transcripts encoding host ribosomes.

## Results

### Human MDMs support productive infection with IAV

After infection of MDMs from four different donors with influenza A/Udorn/72 (H3N2; hereafter, IAV) at a multiplicity of infection (MOI) of 5(Figure S 1 A), some cells were positive for viral antigen (IAV nucleoprotein) by immunofluorescence after 2 hours and the large majority after 7 hours. This suggested that viral mRNA molecules were being transcribed and translated (Figure S 1 B). RNA libraries were prepared from cells 0, 2, 7, and 24 hours post-infection and from two uninfected-infected samples at 0 and 24 hours. Libraries were sequenced using HeliScope CAGE as previously described (Forrest *et al*., 2014; Kanamori-Katayama *et al*., 2011). MDMs were exposed to IAV for 1 hour before the culture medium was replaced with fresh medium lacking IAV therefore the 0 hours post-IAV time point refers to cells harvested after the initial IAV exposure and immediately prior to medium replacement. We confirmed a previous report (Hoeve *et al*., 2012) that IAV-infected MDM cells released infectious virus (Figure S 1 C), albeit at low levels compared to published results for A549 epithelial cells and with little evidence of cell death up to 7 hours (Figure S 1 D). Despite producing relatively little infectious virus, the majority of the IAV-infected MDMs were lost after 24 hours.

### Network analysis of the response to influenza virus infection in MDM

RNA libraries were prepared from similarly infected cells at 0, 2, 7, and 24 hours post-infection and from two uninfected samples at 0 and 24 hours. Libraries were sequenced using HeliScope CAGE as previously described (Forrest *et al*., 2014; Kanamori-Katayama *et al*., 2011). Note that MDMs were exposed to IAV for 1 hour before the culture medium was replaced with fresh medium lacking virus; therefore the 0 hours post-infection time point refers to cells harvested after the initial IAV exposure and immediately prior to medium replacement. In addition, expression profiles of the MDM at 24 hours likely reflect the remaining survivors of the IAV infection, since dead cells were detached from the plates and were thus not lysed for RNA extraction.

### Network analysis of the response to IAV virus infection in MDM

Temporal changes in host cell transcription are likely to occur both in recognition of viral infection and as a consequence of viral lifecycle progression. This study gave us the opportunity to observe changes in transcription at 4 time points over 24 hours of IAV infection. We utilised the network analytical tool, Graphia (Freeman *et al*., 2007), to identify sets of co-regulated transcripts in the MDM response to IAV (Table S 1). For simplicity, we restricted the analysis to the dominant (most frequently used) promoters (p1) and used averaged data from the 4 donors. We have summarised the GO term enrichment and pathway enrichment in the 10 largest clusters using GATHER (Chang and Nevins, 2006) (Table S 2) and Enrichr (Chen *et al*., 2013; Kuleshov *et al*., 2016) (Table S 3) respectively.

Figure 1 A shows the sample-to-sample correlation graph for each of the averaged data sets. Although there was a global alteration in gene expression that progressed with time, the profile at 7 hours remained correlated with the profiles in uninfected cells at both early and late time points. This suggests that the virus did not cause a selective, or global, loss of expression of host-related genes. In keeping with that conclusion, the largest cluster, Cluster 1, contained more than 4,500 genes (Figure 1 B) whose shared pattern was continuous induction across the time course with particularly high expression at 24 hours. This cluster contained genes encoding the interferon-responsive transcription factors, *IRF1, 2, 4, 7, 8, and 9* and numerous known interferon-responsive antiviral effector genes (e.g. *APOBEC3G,* RSAD2, *DDX58, ISG15, MX1*, *OAS1, TRIM25*). Cluster 2, at less than half the size, is the near-reciprocal cluster to Cluster 1 as it contained genes that were progressively downregulated by IAV up to 7 hours post infection. A prominent feature of the GO annotation and pathway analysis of this cluster is protein synthesis, secretion and intracellular transport (Table S 2, S 3), exemplified by multiple Golgi-associated genes (e.g. *GOLGA2, GOLPH3, GGA3*), components of the coatamer complex (*COPA, COPG*) and secretory components.

**Figure 1.**
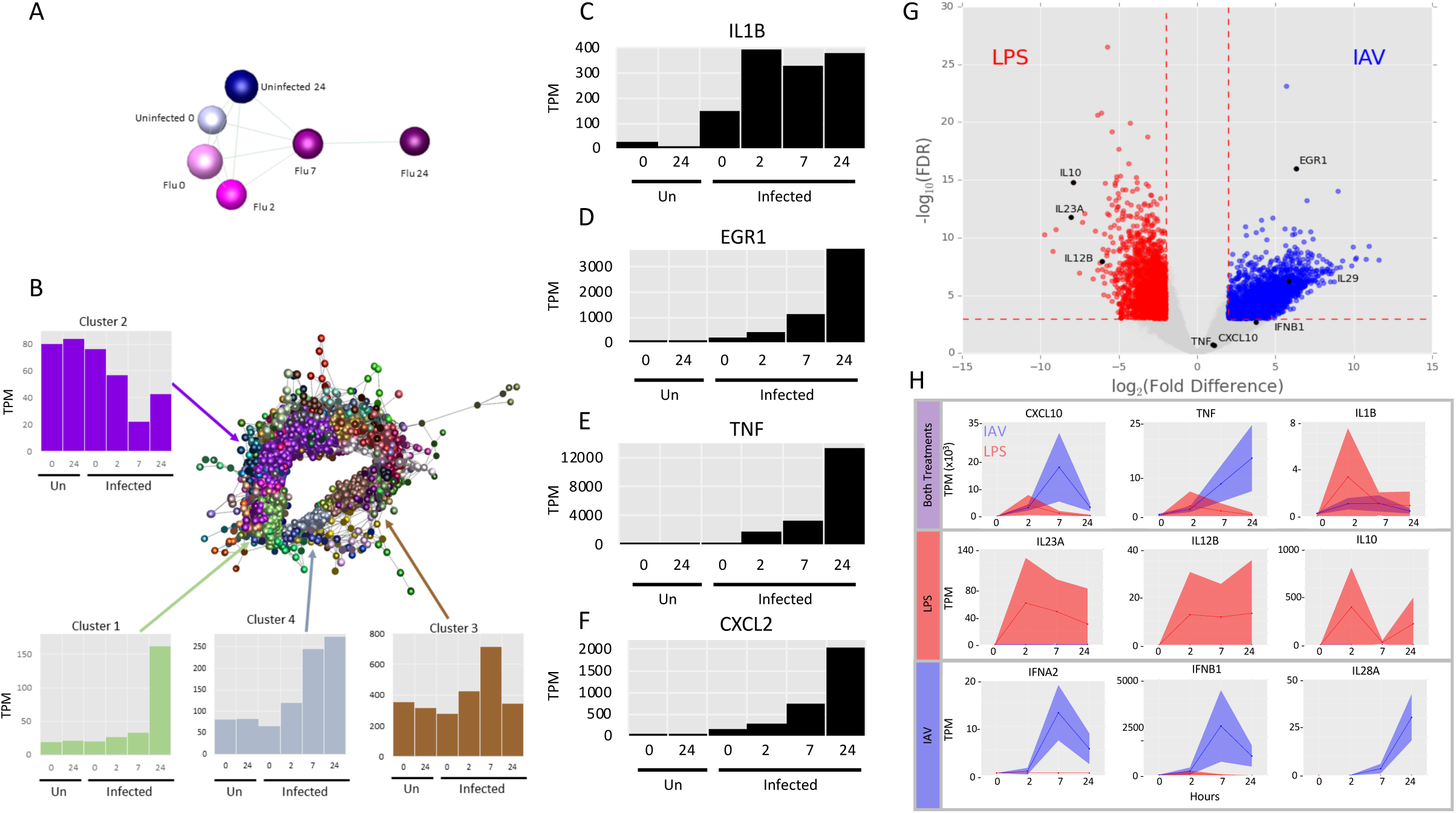
Network analysis of the co-expressed genes during IAV infection in MDMs demonstrates their rapid response. (A) Sample-to-sample network. A correlation coefficient of ≥ 0.7 was used to include all samples in the network. Analysis was restricted to the dominant promoters (p1) and data were averaged across the 4 donors. Blue – uninfected; pink – infected; darker colours show later time points. (B) Gene-to-gene correlation profile of transcripts. Network analysis identified the sets of co-regulated transcripts in the MDM response to IAV. Analysis was restricted to the dominant promoters (p1) and data were averaged across the 4 donors. Lines represent connections at Pearson correlation coefficient ≥ 0.94 and spheres represent genes (promoters). The clustering procedure used a relatively coarse Markov clustering algorithm of 1.7 to avoid excessive cluster fragmentation. The four largest clusters, along with their average expression profiles, are shown. Y axis in the expression profiles shows the expression level in tags per million (TPM). (C-F) Abundance of transcripts for IL1B (C) EGR1 (D), TNFα (E) and CXCL2 (F) at the indicated time points. y-axis shows expression in tags per million (TPM). (G) Differential gene expression analysis comparing expression of transcripts in LPS-treated and IAV-treated monocyte derived MDMs. Transcripts with a relative log fold change (log_2_FC) ≥ 2 and a -log_10_(FDR) ≥ 3 are shown in red (higher in LPS treated) and blue (higher in IAV infection). Genes with greatest difference in expression are labelled. Genes referenced in the text are shown in black. (H) Comparison of the temporal response of genes between IAV- and LPS-treated MDMs. Expression (TPM) of selected genes in LPS-treated (red) and IAV-infected (blue) human MDMs at 0, 2, 7, and 24 hours post treatment is shown in tags per million (TPM). Solid lines show the mean expression of all donors (n = 3 for LPS, n = 4 for IAV). Filled-in area shows standard deviation between donors.

Clusters 3 and 4 had similar profiles to each other, differing only at the late time point of 24 hours, and between them contained a set of rapidly-induced genes, including interferons *IFNB1*, *IFNA1, IFNA2*, *IFNA8, IFNA14*, *IFNE* and further known IFN-regulated targets such as *IFI6, IFIT2, IFITM3*, *IRG1*, *GBP1* and *MNDA*. The enrichment of these clusters also highlights induction of genes involved in protein synthesis, including 46 ribosomal protein subunit genes and those associated with the mitochondria, oxidative phosphorylation and ATP synthesis. We observed that the response of MDMs to viral infection was immediate. IL1B was rapidly and strongly induced by IAV at 0 hours (effectively 1 hour post virus addition) and peaked at 2 hours (3 hours post virus addition) (Figure 1 C). Other early response genes that were detected early after IAV exposure included those encoding immediate early transcription factors such as EGR1, the proinflammatory cytokine TNFα and the neutrophil chemoattractant CXCL2 (Figure 1 D-F).

### Comparative analysis of the response of MDMs to treatment with IAV and with LPS

The response of MDMs to IAV and LPS was compared at equivalent time points, uncovering some common transcripts that were expressed in both treatments (Figure 1 H, top row). Transcripts induced specifically by LPS but not by IAV were revealed by differential expression analysis (Figure 1 G, Table S 4) and included classical inflammatory cytokines IL12B (although not IL12A) and IL6, and the feedback regulator of inflammation, IL10 (Figure 1 H, central row; Figure S 2 A, B). Conversely, induction of genes associated with interferon signalling was more substantial and prolonged in IAV-treated MDM than those treated with LPS. IAV induced *IFNB1* mRNA some 10-fold more than observed in response to LPS in MDM, and sustained this expression throughout the time course (Figure 1 H, bottom row). IAV also induced multiple IFNA genes (*IFNA1, A2, A8, A14, A22*, Figure S 2 C-E) and the type III interferon genes, *IFNL1* (aka IL28A) and *IFNL2* (aka *IL29*), which were not induced at all by LPS (Figure 1 H, bottom row; and Figure S 2 F).

### Transcriptional activity of IAV in human MDMs

IAV mRNAs contain a conserved 12-base long 5’-adjacent non-coding region present in all 8 segments (‘AGCAAAAGCAGG’) derived from template-dependent transcription of the viral promoter (De Vlugt, Sikora and Pelchat, 2018). Possession of this sequence was therefore used to identify viral transcripts. Similar to results seen elsewhere (Gu *et al*., 2015; Koppstein *et al*., 2015; Sikora *et al*., 2014), the A at the 5’ end of the promoter was not always present and thus sequences which contained the 11 nucleotide sequence ‘GCAAAAGCAGG’, referred to subsequently as the IAV promoter, were brought forward for analysis. The IAV promoter is present in all 8 viral mRNA segments and follows the host-derived leader sequence (Figure 2 A).

**Figure 2.**
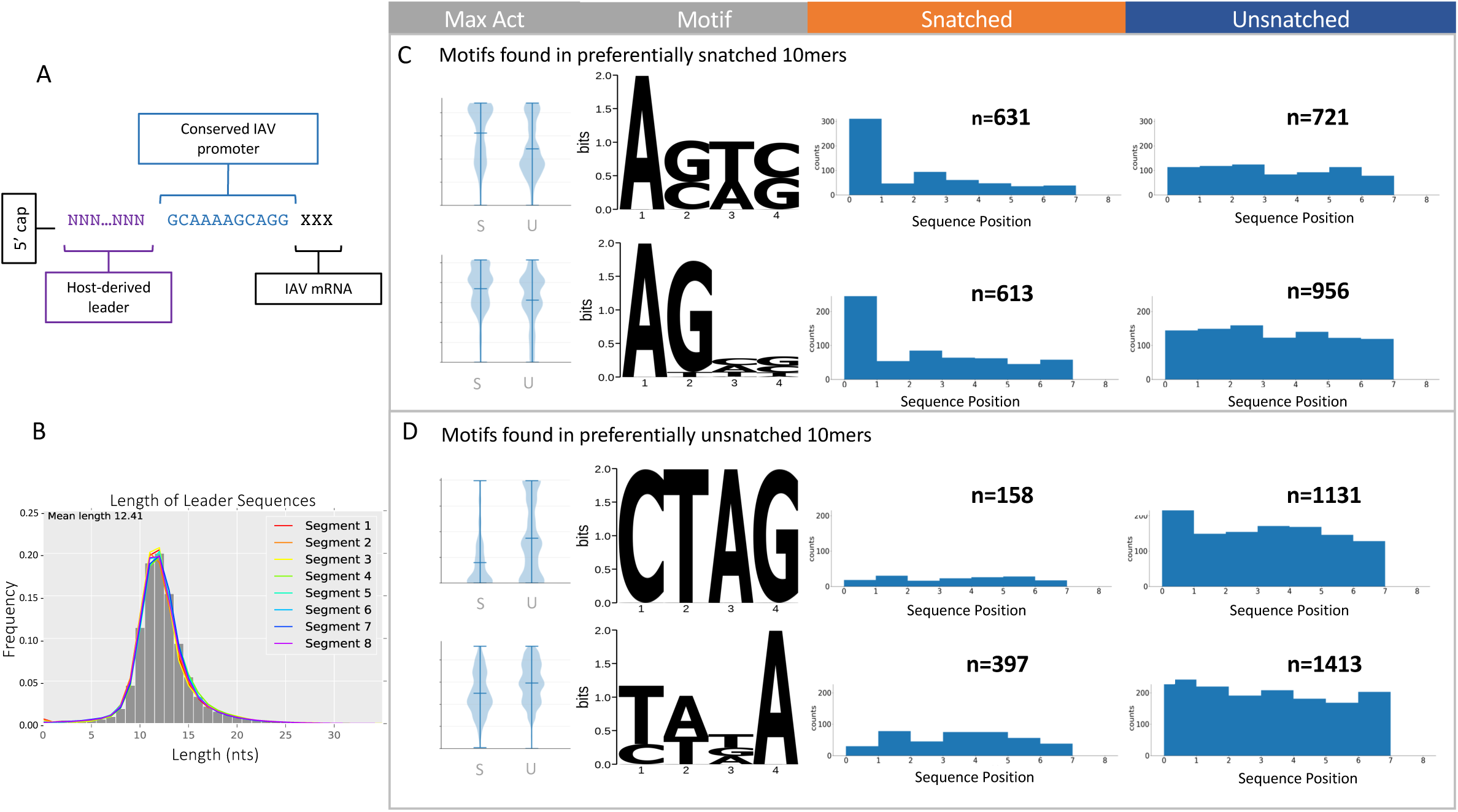
Motifs associated with snatched and unsnatched leader sequences. (A) Schematic showing the structure of the capped 5’ end of IAV mRNAs. (B) Length of leader sequences across segments. Segments are coloured as shown in the legend. (C, D) The first ten nucleotides of each CAGE tag were extracted and the abundance of each sequence associated with IAV was compared to the background abundance by Fisher’s Exact test (FDR < 0.05). Identification of motifs associated with snatched (B) and unsnatched (C) sequences. Violin plots show the maximum activation distributions for snatched (S) and unsnatched (U) sequence categories in arbitrary units. The four-nucleotide long motifs associated with each category are visualised as position weight matrices. The positional enrichment of the four-nucleotide motifs across the 10mer sequences is shown. The number of sequences is given as n above each bar chart.

Overall, the relative proportion of IAV mRNA arising from each viral segment was remarkably consistent across the 4 donors at each time point, demonstrating the coordinated nature of transcription by IAV (Figure S 3 A). Published studies of A549 cells reported that, within 8 hours post-exposure, >50% of total cellular mRNA was viral (Bercovich-Kinori *et al*., 2016). In contrast, in the infected MDM capped IAV RNA constituted a relatively small proportion (4 - 11%) of total capped RNA in the cell even at the peak of viral replication (Figure S 3 B). The relative proportion of IAV mRNA arising from each viral segment was remarkably consistent across the 4 donors at each time point (Figure S 3 C) compatible with the understanding that transcription of each segment is a highly controlled process (McCauley and Mahy, 1983). The relative proportions of the viral transcripts encoding the polymerase segments (1, 2, 3, encoding PB2, PB1, and PA respectively) peaked at the 0 hour post-infection time point (after 1 hour incubation with IAV), together with the detectable induction of host response genes observed above. Relative expression of segment 8 (NS1/NS2) was highest at 2 hours. The late structural segment transcripts (4, 6, 7, encoding HA, NA and M1/M2 respectively) peaked at 7 hours, towards the end of the expected 6-8 hour viral life cycle. By 24 hours, the pattern was less defined, which may be a consequence of mRNA decay or potential reinfection by the virus. In addition, reads plausibly corresponding to the known mRNA3 splice variant transcript from segment 7 which utilises a splice donor site at the 3’-boundary of the conserved promoter sequence (Lamb, Lai and Choppin, 1981) were also seen (Figure S 3 D). Similar sequences that potentially represent alternative splice variant mRNAs were observed, most abundantly from segments 5 and 6 (Figure S 3 D, Table S 5). It is unknown if these can be translated.

### Characterisation of host leader sequences incorporated into viral capped RNA

We identified 4,575,918 unique leader sequences, heterogeneous in both sequence and length, snatched from the host and incorporated into viral mRNA. Of these leader sequences, 18.8% (859,789) appeared more than once and 1.5% (69,443) appeared ten times or more across all samples. This 1.5% of the most frequently occurring accounted for 53.6% of the total number of snatched leaders. Thus, at least two populations of snatched sequences exist: those that were heavily snatched as they occur multiple times, and those that were seemingly randomly snatched as they appear only once. Most (74%) of the leader sequences were between 10 and 14 nucleotides long (Figure 2 B). Contrary to previous reports (Koppstein *et al*., 2015; Sikora *et al*., 2017), we observed no difference in leader lengths among different viral segments.

We sought to determine whether there was over-representation of particular sequences, host transcripts, or biological pathways among the total population of leader sequences. In order to eliminate the risk of bias due to the different rates of successful mapping for sequences of different lengths, we restricted our analysis to the first 10 bases of every CAGE tag (10mers) meeting the abundance threshold of 1,000 tags across all examined samples in the dataset, including both IAV and host sequences. The number of times a 10mer was followed by the IAV promoter, i.e. incorporated into viral mRNA (“snatched”), was compared to the number of times a 10mer was not followed by an IAV promoter (“unsnatched”) using Fisher’s Exact test (FDR <0.05) at each time point (see Methods). We uncovered patterns of significant over- and under-representation for specific 10mers.

### Enrichment of specific RNA motifs in the snatched and unsnatched sequence populations

It is not known if the cap-snatching mechanism targets specific nucleic acid sequences. However, leader sequences are known to commonly have ‘GCA’ at the interface between the host sequence and the IAV promoter (Rao, Yuan and Krug, 2003; Geerts-Dimitriadou, Goldbach and Kormelink, 2011) partially as a consequence of the “prime and realign” mechanism of IAV mRNA transcription (Beaton and Krug, 1981; Geerts-Dimitriadou *et al*., 2011; Koppstein *et al*., 2015). More recently, an ‘AG’ at the 5’ end of the leader sequence has also been shown to be prevalent in snatched sequences (Gu *et al*., 2015).

Our analysis of 10mers enables a statistically powerful comparison of snatched and unsnatched sequences in which the position of sequence motifs can be compared without reference to distance from the 5’ or 3’ ends. We used Pysster (Budach and Marsico, 2018) to train convolutional neural networks using the sequence data to explore sequence and positional features for pools of highly-significantly over-represented snatched and unsnatched 10mers (0.3 >= OR >= 3, -log(FDR < 10). This stringency was introduced to eliminate potential noise. Optimisation experiments indicated that a kernel (sequence motif) length of 4 had relatively consistent recall and high precision for this dataset (Figure S 4). The snatched 10mers showed an enrichment of two motifs, A[G/C][T/A][C/G] and the similar sequence AGNN, both beginning at the first base (position 0) (Figure 2 C). These motifs were most apparent 2 hours post infection, coinciding with levels of high transcription by the virus and are consistent with previous reports of an ‘AG’ preference at the 5’ end of the leader (Gu *et al*., 2015).

The unsnatched 10mers also showed an enrichment of two distinct motifs, CTAG and [T/C][A/T][T/G/A]A, most evident at 7 hours post infection (Figure 2 D). While the CTAG motif was unsnatched primarily when it began in the first position (position 0), there was also an association between this motif at any position in the 10mer and unsnatched status. Similarly, the [T/C][A/T][T/G/A]A motif was preferentially avoided by cap snatching if it occurred at any position within the 10mer. To our knowledge this is the first evidence for the avoidance of particular sequences as priming leaders by the IAV polymerase.

### Host genomic origin of over- and under-represented sequences

Of 29,195 10mers, we assigned transcript identity to 12,992 (44.5%)(Methods) and of the named 10mers, 6,353 mapped to more than one transcript; for these, a single transcript was chosen at random from the list of possible sites. 8,895 10mers had annotated host promoter/gene names and did not contain IAV promoter-like sequences (Methods). The remainder are a mixture of alternative host promoters, lncRNAs, eRNAs and other RNA species (Andersson *et al*., 2014). This approach costs statistical power, but is necessary to avoid any bias that might be introduced into the identification based on a quantitative measure, such as abundance. We repeated the Fisher’s Exact analysis for all 10mers assigned to a gene name to provide summary data for that gene. The 1,000 most significantly enriched named genes in the snatched and not snatched sets are reported in Table S 6.

A cap-snatching event can occur at any point after infection and before and RNA extraction from the cells, therefore a more relevant background pool of host RNAs from which a given leader could have been obtained may be the host RNA content at the preceding time point. Our dataset allowed us to systematically compare every 10mer in infected cells against the background RNA in the cell from the same time point, and also against the background RNA at a previous time point. Genes that met this stringent threshold, for at least two donors at all time points, are reported with a description of their function in Table S 6.

### Host transcriptional machinery is potentially targeted by the cap-snatching mechanism

Key spliceosome snRNAs (*RNU1, RNU11, RNU12, RNU4ATAC, RNU5A, RNU5E, RNU5F, RNU5D, RNU7*) and their variants/pseudogenes were high among the most significantly enriched named genes. This is consistent with previous observations that snRNAs are snatched frequently and shows that this may represent a true preference for these RNAs. In view of the preferential snatching of multiple snRNAs, we considered whether specific classes of capped host RNAs might be targeted. Of the RNA types we considered, only snRNAs were strongly preferentially snatched (Figure S 5 A, B).

This sequencing method also allows the observation of histone mRNA which enabled us to observe that 10mers corresponding to histone mRNAs were also significantly over-represented. In addition, we observed that many host mRNAs encoding spliceosome- and transcription-associated proteins (*SRSF3, SRSF6, PRPF18, SNRNP25, SNRNP70, MAGOH*) were preferentially snatched. The 10mer corresponding to the transcript encoding the largest subunit of RNA polymerase II (POLR2A), was 5.83-fold over-represented in snatched sequences (OR = 5.83; FDR <0.05). *PABPN1*, which encodes poly(A) binding protein, was also preferentially snatched (OR = 2.28; FDR <0.05). These comprise key elements of both transcription and polyadenylation of host mRNAs. Taken together these observations imply that cap-snatching may interfere with regulation of transcription and splicing in the infected cell. However, POLR2B, another subunit of RNA polymerase II, was 7.77-fold under-represented (OR = 0.13; FDR < 0.05) making it difficult to draw simple conclusions. To rectify this, we performed gene set enrichment analysis to statistically determine over- and under-represented pathways affected by the cap-snatching mechanism.

### Pathway enrichment analysis indicates that specific ribosome-associated transcripts are avoided by the cap-snatching mechanism

Identified transcripts from all time points and donors were collated and gene set enrichment analysis performed by querying various pathway/gene ontology datasets (listed in Table S 7). Querying Reactome gave a single over-represented pathway: RNA Polymerase II transcribes snRNA genes (Figure 3 A). A volcano plot highlighting the distribution of pathway members shows that many mRNA pathway members were under-represented and the members of this pathway that drive its enrichment as an over-represented pathway were predominantly snRNA transcripts (Figure 3 B), particularly snRNA members of the minor spliceosome. This is consistent with the observed preferential snatching of snRNAs. These transcripts were upregulated in IAV treated MDMs compared to LPS treated MDMs (Figure S 5 C).

**Figure 3:**
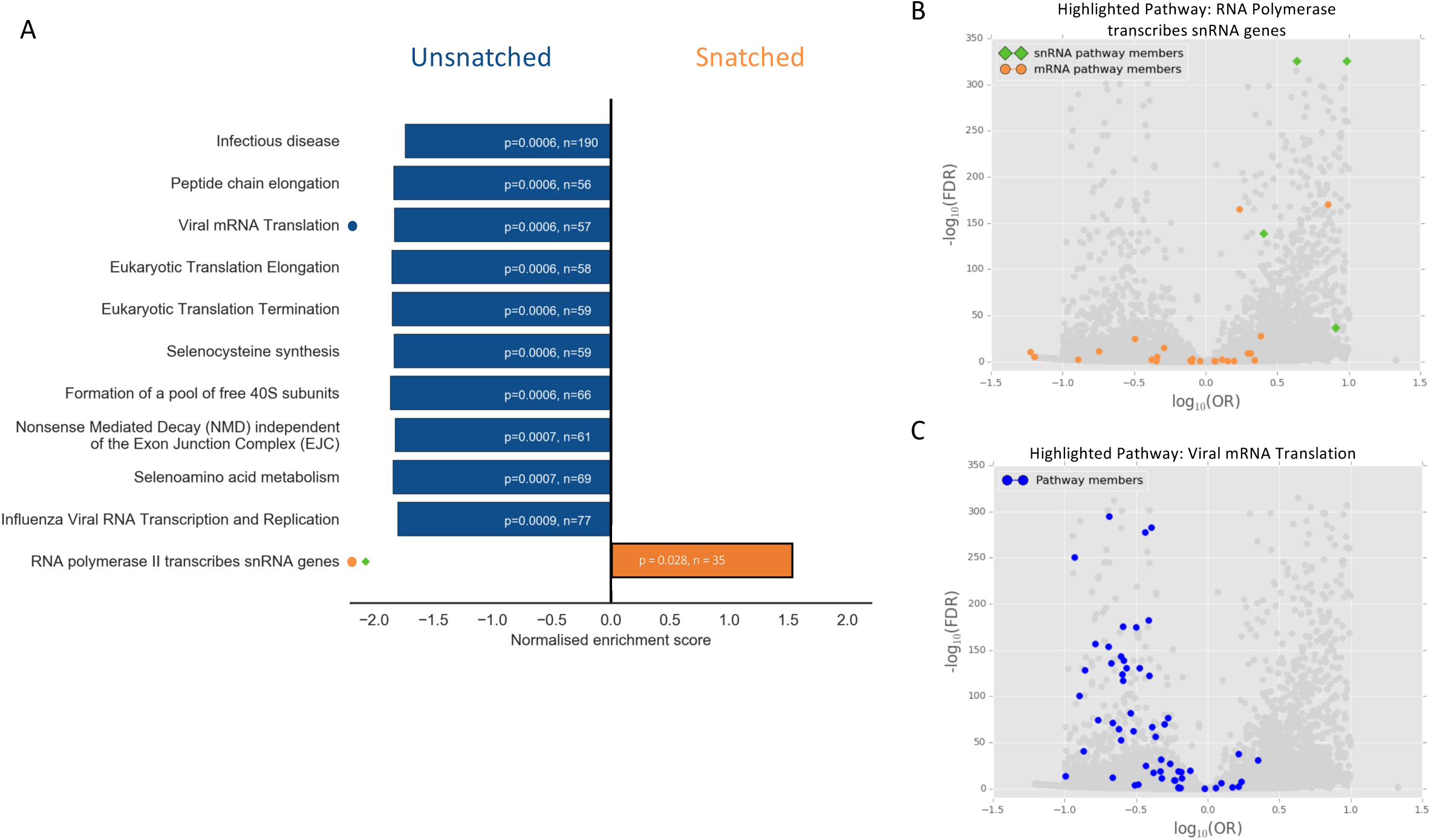
Pathways that were enriched in snatched and unsnatched sequences. (A) The 10 most under-represented pathways (negative enrichment score, blue) and single significantly over-represented pathway (positive enrichment score, orange) in the Reactome 2016 database are shown. N represents the number of genes associated with that pathway detectable in the dataset. p-values shown are Benjamini-Hochberg FDR-adjusted p-values. (B) Volcano plot showing the significance as -log_10_(FDR) and odds ratio of snatched versus unsnatched 10mers with members of the Reactome pathway ‘RNA Polymerase transcribes snRNA genes’ highlighted (snRNA, green diamonds, mRNA orange circles). (C) The same volcano plot as in (B) with members of the Reactome pathway ‘Viral mRNA Translation’ highlighted (blue circles).

Pathway enrichment also allowed us to look for pathways that were avoided by the cap-snatching mechanism. We identified pathways associated with translation and ribosome formation as significantly under-represented in the cap-snatched pool (Figure 3 C). Although multiple pathways were identified, these were not independent: these associations were largely driven by presence of a group of transcripts encoding the same set of ribosomal proteins (Table S 7). These data show that IAV avoids snatching caps from ribosomal mRNA transcripts. Interestingly, not all mRNAs encoding ribosomal subunits were avoided. We compared our results to a recent study reporting the effect of targeted knockdown of specific ribosomal subunit mRNAs in the context of IAV infection (Wei *et al*., 2019), but saw no clear relationship between cap-snatching preference and viral protein production, host protein production, or antigen presentation.

The 10mers associated with the ribosomal protein transcripts in question, and their pseudogenes, which are generally indistinguishable at the 10mer level, were extracted and aligned using Meme (Bailey and Elkan, 1994). Amongst the analysed 10mers the most commonly observed motif was CTCTT[T/C]C[T/C] (p < 0.05) (Figure S 5 D) originating at the first position. This is broadly in agreement with our observation above that CTAG occurring at various positions within the 10mer is unlikely to be snatched by IAV, regardless of abundance.

## Discussion

This comprehensive analysis of host and viral transcripts reveals key features of host-pathogen interaction at a molecular level. We demonstrate that IAV cap-snatching has a strong preference for host transcripts associated with splicing and transcription, and avoids host ribosomal subunit transcripts. By comparing against a canonical innate immune stimulus, LPS, we systematically characterise the host response of human macrophages to IAV exposure.

### Characteristics of MDM response to IAV

MDMs can be infected with IAV and produce both viral protein (NP) and infectious virus. This initial permissiveness may be related to the fact that *IFITM3*, the protein product of which restricts viral infection and is associated with IAV susceptibility (Everitt *et al*., 2012), was almost completely down-regulated in MDM compared to the high level of expression in blood monocytes (can be observed at http://fantom.gsc.riken.jp/zenbu/).

Wang *et al*. (Wang *et al*., 2012) discussed the possible cellular pattern receptors required for recognition and response to IAV infection in MDMs. Of those candidates, mRNAs encoding RIG-I (DDX58), MDA-5 (IFIH1) and TLR3 were expressed at very low levels in MDMs. In contrast, TLR7 was expressed at similar levels in MDM to the level in plasmacytoid dendritic cells (http://fantom.gsc.riken.jp/zenbu/) and thus appears the most likely initial intracellular receptor that initiates the response to viral nucleic acid.

Examination of co-expression clusters suggests that in MDMs IAV does not cause a selective or global loss of transcription of host-related genes. The GO enrichment for the largest cluster observed, Cluster 1 (Table S 2), included the ubiquitin-proteasome complex, oxidative phosphorylation, cell cycle and transcriptional regulation including mRNA splicing and binding. Together with the consistent similarity in global gene expression between uninfected and early post-infection time point, this suggests that most basic cellular processes are maintained during infection. In A549 cells, IAV infection causes cell cycle arrest (He *et al*., 2010), and down-regulation of cell-cycle associated genes. Since MDM are not actively proliferative, the apparent induction by IAV infection of many cell cycle-related genes, including the cyclin genes *CCNA1, CCNB1, CCND1, CCNE1, CCNE2* and *CCNG2* and 19 genes encoding multiple cyclin-dependent kinases (CDK) is unlikely to be associated with cellular proliferation.

The gene set induced by IAV infection likely includes many additional anti-viral effectors. For example, the gene encoding BST2-tetherin, which inhibits the release of enveloped virus particles, possibly including some strains of IAV (Gnirß *et al*., 2015) was induced by both LPS and IAV. A neighbouring gene, *MBV12A*, shares a bidirectional promoter with *BST2,* a subunit of the ESCRT complex, over-expression of which interferes with viral assembly in HIV-1 infected cells (Morita *et al*., 2007). *MBV12A* was much more strongly induced by IAV than *BST2* in MDM, and was only marginally induced by LPS in the same cells. Similarly, expression of the antiviral effector *RSAD2* (viperin) (Wang, Hinson and Cresswell, 2007; Gizzi *et al*., 2018) was induced at 2 and 7 hours. Many other inducible genes in Cluster 1, including multiple members of the *FOX, TRIM, CDK, USP* and *MED* families have been shown to be phosphorylated during the MDM in response to IAV (Söderholm *et al*., 2016) and are implicated as antiviral effectors (Nyman *et al*., 2000; Ohman *et al*., 2009).

A feature of Cluster 2 that is not evident from GO annotation, is the ablation within 7 hours of transcripts encoding many cell surface receptors and signalling molecules with roles in innate immunity, including *CSF1R, C3AR1, C5AR1, CD4, CD14, MYD88, CD180, CD44, CD163, FCGR2A, IL10RB, ITGAM, ITGAX, TLR1, TLR4, TLR8, TNFRSF1A,* and *TGFBR1*.

### Variation in host transcript expression between donors

Substantial variation amongst MDMs from different individuals was evident in most IFN-inducible genes. Fairfax *et al*. (Fairfax *et al*., 2014) reported that up to 80% of inducible genes in human monocytes responding inflammatory stimuli show evidence of heritable variation in their level of expression. Since genetic variation between hosts alters risk of death from influenza (Horby *et al*., 2012), and variants underlying inter-individual variation in IAV-induced gene expression are associated with human disease phenotypes (Lee *et al*., 2014), we quantified variation in inducible expression among the four donors in this study. The coefficient of variation for named genes at each time point is provided in Table S 8. The antiviral *IFITM3* and the neighbouring *IFITM1* were selectively inducible in Donor 3, albeit to low levels (Figure S 6 A, B). Fairfax *et al*. also found trans-acting expression variants, likely involving autocrine IFNB1 signalling upstream of separate regulons targeted by the transcription factors IRF7 and IRF9 (Hume and Freeman, 2014). One of the four donors in our study, Donor 1 showed a more rapid induction of *IFNB1* in response to IAV, which preceded high *IRF7* and *IRF9* induction (Figure S 6 C, D). Conversely, cells from Donor 4 had a relatively low *IFNB1, IFNA1* and *IRF9* induction in response to IAV, whereas *IRF3* was relatively stable throughout infection in all four donors (Figure S 6 E-G).

### The comparison between the host response in IAV and LPS treated MDMs

Like LPS, IAV strongly induced *TNFα*, *IL1B*, multiple chemokine genes (e.g. *CCL2, CCL3, CXCL1, CXCL2, CCL20*) and many genes for immediate early transcription factors (e.g. *EGR* family, *FOS* family, *JUN* family, *NR4A1, ATF3* etc.). However, the global gene-based analysis of the response of MDM to IAV reveals a clear contrast to the LPS response in MDMs. In LPS-treated human MDMs mRNA levels of proinflammatory genes are subject to control by a complex network of rapidly-inducible feedback regulators including *DUSP1, TNFAIP3, NFKBIA, ZC3H12C, PTGS2* and the microRNA *mIR-155* (Baillie *et al*., 2017). The sustained induction of proinflammatory transcripts in response to IAV contrasts with this transient induction in response to LPS. Each of these feedback regulators was induced to a lesser extent and/or much later in the response, by IAV compared to LPS.

Unlike Hoeve *et al*, (Hoeve *et al*., 2012) we did not detect expression of transcripts encoding either of the subunits of IL12 (*IL12A, IL12B*), prior to the 24 hour sample, in MDM in response to IAV. Following LPS treatment, MDM have low expression of *IL12A* (p35) (Figure S 2 B), instead inducing *IL23A* and *IL12B* mRNA, which together encode the heterodimeric proinflammatory cytokine IL23. These were not detected in IAV-infected cells. Similarly, there was no detectable induction of the anti-inflammatory cytokine *IL10* mRNA by IAV, in contrast to the massive and sustained induction by LPS.

The type III interferons were highly specific to IAV-treated MDMs. These were recently shown to mediate a key mechanism preventing viral spread to the lower respiratory tract in mice (Klinkhammer *et al*., 2018), which is believed to cause life-threatening disease in humans (Van Riel *et al*., 2007). The sustained induction of IFN responsive genes (Cluster 1, see above) shows that induction of IFN and IFN signalling is clearly not successfully prevented by the primary IAV interferon antagonist NS1 in MDM, by contrast to the pattern observed in other cell types (Haye *et al*., 2009; Jia *et al*., 2010; Thakar *et al*., 2013; Perez-Cidoncha *et al*., 2014). The observed constraint on production of new virus may be attributed to the massive interferon response (Cluster 1, see above), and down-regulation of synthesis of secreted proteins (Cluster 2, see above). This profound difference in induction of IFN-responsive genes between LPS and IAV stimulation is reflected in blood transcriptome profiles of patients with severe IAV compared to those with bacterial sepsis (Ramilo *et al*., 2018).

### Elimination of bias for accurate quantification of leader sequences and 5’ RNA ends

Our choice of sequencing methodology and analytical approach eliminated numerous sources of bias that have limited the interpretation of previous studies of cap-snatching preference. A key difference from previous work is the accurate quantification of background transcription, which enables the first accurate quantification of the transcripts *not* snatched by IAV.

The HeliScope single molecule CAGE sequencing methodology has several key advantages. This method sequences transcripts from the 5’ end without internal segment-specific primers, and without PCR amplification (Kanamori-Katayama *et al*., 2011). In contrast, previous studies of IAV virus transcripts used internal primers for the viral segments (Koppstein *et al*., 2015; Sikora *et al*., 2017) or performed library amplification on cDNA derived from capped RNA (Gu *et al*., 2015).

In addition, our use of terminal-depth sequencing limits noise and sampling error, in both the snatched sequences and the background distribution. Since CAGE reads sequences directly from the 5’ end, we can be confident that we have quantified the background pool of potential leader sequences that were available to be snatched. By limiting our analysis to sequences of a specific length (10mers), we eliminate bias that may occur due to differential mapping or identification of sequences of different lengths.

Our timecourse design allows us to mitigate another potential source of bias. An unknown period of time has passed between a cap-snatching event and RNA extraction from the cells. Therefore, the relevant background pool of host RNAs from which a given leader could have been obtained is the host RNA content at a previous time. We systematically compared every enriched sequence in infected cells against the background RNA in the cell from the same time point, and against the background RNA at a previous time point (Table S 6).

### Host mRNA processing machinery is preferentially targeted by IAV cap-snatching

Non-coding RNAs, particularly snRNAs, have been identified as the source of the most frequently snatched leader sequences (Gu *et al*., 2015; Koppstein *et al*., 2015). However, it was unclear whether this frequency reflected their relatively high abundance or true over-representation of this RNA type among leaders. Our use of terminal-depth sequencing of complete 5’ sequences, combined with our focused analysis on 10mers, enables an unbiased, accurate quantification of the abundance of each sequence in both the snatched, and unsnatched, sequence sets.

Differential expression analysis revealed that all snRNAs, apart from RNU1, were upregulated in IAV treated MDMs compared to LPS (Figure S 5 C). Notably, snRNA components of the minor spliceosome (*RNU11, RNU12, RNU4ATAC, RNU5A* and *RNU5E*) were highly preferentially snatched, particularly at 2 and 7 hours. In the FANTOM5 dataset, the components of the minor spliceosome were most expressed in the later time points of MDM infection with IAV (can be viewed in Zenbu Browser). *RNU6ATAC* is the only snRNA component of the minor spliceosome we did not observe to be snatched, despite upregulation during IAV infection. This mRNA is transcribed by RNA polymerase III (Singh and Reddy, 2006; Canella *et al*., 2010) leading to a different cap structure which may not be recognised by the IAV polymerase (Koppstein *et al*., 2015). The minor spliceosome splices <1% of introns in the human genome and its activity – and hence functional expression of these splice variants – is regulated by *RNU6ATAC* and increased by signalling through the p38MAP kinase pathway (Younis *et al*., 2013). The viral NS1 protein is known to inhibit the formation of *RNU12/RNU6ATAC* complexes (Wang and Krug, 1998). Our results suggest that IAV may have evolved more than one mechanism to suppress gene expression through the minor spliceosome pathway.

### RNA that codes for ribosomal subunits is avoided by IAV cap-snatching

Our comparison of LPS and IAV-treated cells shows that genes encoding ribosomal subunits are highly differentially transcribed in IAV-treated cells. Therefore, if cap-snatching were primarily determined by abundance, as previously thought (Sikora *et al*., 2017; De Vlugt, Sikora and Pelchat, 2018), we would expect to see leader sequences derived from ribosomal genes prominently among the snatched sequences. Explorations of leader sequence analysis have focused on the snatched population of sequences out of necessity. Our analysis allowed us to determine those sequences that remained unsnatched in the host cell. Although we do see a minority of ribosomal protein mRNA snatched, IAV cap-snatching exhibited a surprisingly strong avoidance of mRNAs encoding ribosomal proteins, which is particularly evident in our pathway enrichment analysis. Recent evidence shows that altering the relative abundance of particular protein subunits of the ribosome can specifically affect the presentation of IAV encoded proteins by MHC-I (Wei *et al*., 2019). This suggests the hypothesis that the virus has evolved to avoid inadvertently altering ribosome abundance and/or composition in a manner would be deleterious to its own replication.

### Limitations of this study

Our study was limited to one cell type and one strain of IAV. It is to our knowledge the most comprehensive systems-level evaluation of both host and viral transcriptional activity for IAV replication, and the first study to perform an unbiased quantification of cap-snatching preference compared with accurate measurement of background transcription. It is possible that the observed enrichment of capped snRNA may be specific to MDMs. Given the high prevalence of snRNA in the IAV leader sequence population of H1N1 infected A549 cells in other studies (David Koppstein, Ashour and Bartel, 2015; Gu, Glen R. Gallagher, *et al*., 2015), it is reasonable to speculate that this mechanism is generalizable across IAV-infected cell types. Future work is needed to explore the mechanisms underlying the preference and avoidance of specific mRNAs, and to determine cap-snatching preferences of other IAV strains and in other cell types.

## Conclusions

Our combined analysis of host and viral transcriptomes of IAV-infected human MDMs and comparison with the response to LPS reveals IAV-specific features of the host response. In overview, we suggest that MDMs contribute to host defence against IAV in multiple ways. Firstly, MDMs internalise virus and produce viral proteins, but they undergo cell death without generating large numbers of progeny virus. These activities lead directly to viral clearance and since MDMs are professional antigen-presenting cells, also promote presentation of viral antigens to T cells. Secondly, in response to IAV, MDMs generate sustained high levels of multiple interferons providing protection against infection of neighbouring cells, including incoming inflammatory cells. On the other hand, the inability of IAV to induce feedback regulators such as IL10 was associated with induction of proinflammatory cytokines that was transient in LPS-stimulated cells but sustained in IAV-infected cells. This failure of feedback regulation may contribute to pulmonary inflammation that is a feature of severe IAV pathology in a clinical setting (Teijaro, 2014). Finally, many genes encoding surface receptors and signalling molecules required for recognition and response to bacteria such as *CD14* and *TLR4* were down-regulated during IAV infection, which may increase susceptibility to secondary bacterial infections that produce morbidity and mortality in IAV infected patients (Morris, Cleary and Clarke, 2017).

HeliScope CAGE at terminal depth allows for an unbiased observation of both IAV segment mRNA and host-derived leader sequences in cap-snatching. Our comprehensive analysis of leader sequences identified motifs that the IAV polymerase may favour and two others that it apparently avoids during the snatching process. We discovered strongly preferential cap-snatching of host sequences associated with splicing, with evidence of the avoidance of key cellular components required for viral replication. These results hint at a mechanism of host evasion through which IAV may downregulate RNA processing machinery through cap-snatching while specifically evading altering translational machinery specifically required for the replication of the virus.

## Materials and Methods

### Ethics, cell culture, virus propagation and infections

Cells were isolated from fresh blood of volunteer donors under ethical approval from Lothian Research Ethics Committee (11/AL/0168). Primary CD14^+^ human monocytes were isolated from whole blood as described previously (Irvine *et al*., 2009) from 4 human donors. Monocytes were plated for 7 days in RPMI-1640 supplemented with 10% (vol/vol) FBS, 2 mM glutamine, 100 U/ml penicillin, 100 μg/ml streptomycin (Sigma Co.), and 10^4^ U/ml (100 ng/ml) recombinant human colony-stimulating factor 1 (rhCSF1; a gift from Chiron, Emeryville, CA, USA) for differentiation into macrophages. Cells were maintained at 37°C with 5% CO_2_. A/Udorn/72 (H3N2) was generated as described previously (Hoeve *et al*., 2012). Differentiated macrophages were infected on day 8. Cells were washed in serum free media after which they were infected at MOI 5 in a volume of 200μl infection media. Cells were incubated for 1 hour at 37°C then washed three times with serum-free media and incubated in RPMI-1640 supplemented with 1μg/ml TPCK-trypsin, 0.7% BSA, 2mM glutamine, 100 U/ml penicillin, 100 μg/ml streptomycin (Sigma Co.), and 10^4^ U/ml (100 ng/ml) rhCSF1. Samples were collected at 4 time points post infection/media change: 0 hour (1 hour after addition of the virus), 2 hours, 7 hours and 24 hours. Uninfected samples were also collected at 0 and 24 hours. LPS treatments were carried out as described previously (Baillie *et al*., 2017). Briefly, cells were treated with 10ng/ml bacterial lipopolysaccharide (LPS) from salmonella Minnesota R595 and harvested at time points from 15 minutes to 48 hours after treatment. Only time points with corresponding IAV treated time points were used in this analysis.

### CAGE

RNA was extracted using the Qiagen miRNeasy mini kit (217004). RNA quality was assessed and CAGE was performed as described previously (Takahashi *et al*., 2012) as part of the FANTOM5 project. Virus genome information is available in Table S 9.

### Data Analysis

Computational analysis was performed using custom Python scripts and as described previously (Forrest *et al*., 2014). Custom Python scripts are available at: https://github.com/baillielab/influenza_cage.

### Network Analysis of the MDM transcriptome during infection

Network analysis of the MDM transcriptome during infection was carried out using Graphia Professional (Kajeka Ltd., United Kingdom; http://www.kajeka.com) -formerly Biolayout *Express*^3D^. Results were filtered to exclude any transcript where the maximum value across all samples did not reach 10 tags per million (TPM). The sample-to-sample analysis was performed at a Pearson correlation coefficient of ≥ 0.70. The gene-to-gene analysis was performed at a Pearson correlation coefficient of ≥ 0.94 and used a relatively coarse Markov cluster algorithm inflation value of 1.7 to avoid excessive cluster fragmentation. We restricted the analysis to the dominant promoters (p1) and used averaged data from the 4 donors.

### EdgeR analysis of LPS treated versus IAV treated samples

Differential expression between groups of genes was analysed using the EdgeR package (Robinson, McCarthy and Smyth, 2009) in R version 3.5.1. CAGE data for LPS and IAV datasets were processed as described previously (Baillie *et al*., 2017). Briefly, for each treatment, the expression value for each clustered transcription start site (CTSS) was compared to the expression values of the corresponding time points from all other donors. Values deviating >3SD from the mean of this pool were replaced with the average of the pool. An average expression value for each CTSS from all donors was then calculated. CTSS with a minimum expression level of 10 tags per million in at least one comparable time point, and with a coefficient of variation > 0.5, were included in expression analysis. Samples corresponding to 7 hours post-treatments were carried forward for analysis. We used the glmFit function to fit the models and glmLRT to perform testing between the LPS and IAV treated samples. Benjamini-Hochberg correction was applied to p-values. A significance threshold of FDR < 0.05 was used.

### Identification and analysis of IAV mRNA

Capped IAV RNAs were identified by the conserved 11 base promoter sequence expected to be in all viral mRNA (‘GCAAAAGCAGG’), as described in the text. Sequences that contained the promoter were classified as capped viral mRNA and aligned to the Udorn genome (Table S 9) using custom Python scripts.

### Unbiased analysis of leader sequence preference

The first ten nucleotides of each CAGE tag (10mers) that reached the abundance threshold in our dataset were extracted and this set of unique 10mers were used in subsequent analysis. The abundance threshold was set to 1,000 occurrences across all samples. To determine the 10mer sequences that were over- and under-represented in the snatched population based on background abundance, the number of times a 10mer was associated with the IAV promoter was counted (“snatched”) along with the number of times the 10mer occurred without the promoter (“unsnatched”). These were analysed using Fisher’s Exact test. Benjamini-Hochberg correction was applied to p-values. Significance was determined by an FDR < 0.05. The number of times a 10mer was snatched was compared to the number of times it occurred unsnatched at the previous time point by Fisher’s Exact test.

### Analysis of leader motifs using convolutional neural networks

A sub-set of 10mers that reached the following threshold: 0.3 < (OR) > 3, -log(FDR < 10) were brought forward for analysis of motif preference using convolutional neural networks. We optimised an existing network (Budach and Marsico, 2018) for our use by using altering the parameters to find suitable setting by using the grid search to explore various kernel lengths (2, 3, 4, and 5) and drop rate (0, 0.1, and 0.5); for other parameters, we used the default settings of Pysster (kernel number: 20, convolutional layer number: 2) apart from learning rate at 0.0001 and patience, stopping at 100. Since our analysis was restricted to 10mers, we did not use the pooling method. We randomly selected the training set and validation set in the proportion of 60% and 30% independently. The purpose of this experiment was to explore the existing data, not to make predictions, so we reused the training data to explore the result. Optimisation experiments demonstrated that a kernel length of 4 gave us relatively high, and relatively consistent, precision and recall. We maximised the area under the receiver operator characteristic (ROC) and the area under the precision recall curve. Motifs were considered if they reached a score of at least 50% the maximum score for that time point.

### Assignment of transcript identity to 10mer sequences

CAGE tags were mapped to the human reference genome (hg19) as described (Forrest *et al*., 2014). To identify the possible transcription start site from which a 10mer arose, we extracted every possible chromosomal location for a 10mer that met the abundance threshold of 1000 across all samples from the original alignment BAMfiles created as part of the Fantom5 project. 10mers containing a 6mer from within the IAV promoter (‘GCAAAA’, ‘CAAAAG’, ‘AAAAGC’, ‘AAAGCA’, ‘AAGCAG’, ‘AGCAGG’) were removed. Reference transcription start sites were downloaded from Fantom5. Promoter identity was assigned first using BEDtools 2.25.0 with a window of +/- 5 bases and exact strand match only. For each possible promoter identity the 10mer was mapped to the genomic sequence with a window of +/- 5 bases directly surrounding the coordinates of the assigned transcription start site and exact matches only were used to assign promoter identity.

In order to avoid any effect of abundance that may bias transcript identification, for 10mers with more than one possible promoter identity, a site was chosen at random from the list of possible sites. Promoter names were converted to HGNC format. To determine over- and under-representation of promoters and genes, all 10mers that were assigned to that promoter or gene name were counted and the Fisher’s Exact test was performed. Benjamini-Hochberg FDRs were calculated using the scipy.stats v 0.18.1 statsmodels.stats.multitest.mutlipletests function with method = ‘fdr_bh’. Significance was determined by an FDR < 0.05. RNA type was assigned to gene names using reference data downloaded from Biomart (http://www.ensembl.org). Only named transcripts were assigned an RNA type.

In order to determine if a gene was significantly snatched compared to its abundance at the previous time point, we compared snatched at t to unsnatched at t-1 for the collated values of all 10mers that were assigned that gene name in each sample separately. This was performed for 2hrs versus 0hrs, 7hrs versus 2hrs, and 24hrs versus 7hrs. A gene was declared ‘True’ if the combined p-value for that gene was significant in 2 out 4 donors at that time point.

### Pathway and Gene Set Enrichment Analysis

All named genes that appeared significant were included in this analysis. Gene names were converted to HGNC format for consistency with gene set libraries, excluding unannotated peaks and names with no HGNC equivalent. GO term assignment and pathway analysis for coexpression clusters were performed using Enrichr (<u>mp.pharm.mssm.edu/Enrichr</u>) (Chen *et al*., 2013; Kuleshov *et al*., 2016) and GATHER (Chang and Nevins, 2006). Pathway databases queried were: Reactome 2016, KEGG 2016, WikiPathways 2016 and GO Molecular Function 2015, GO Cellular Component 2015 and GO Biological Process 2015. Gene Set Enrichment analysis on ranked cap-snatching preference data was performed using R package FGSEA (Sergushichev, 2016), in R version 3.5.1, with the following parameters: set.seed = 42, min set size = 5, max size = 5000, nproc = 1, nperm = 1000000. Gene set libraries KEGG 2016, BioCarta 2016, Reactome 2016, WikiPathways 2016, NCI Nature 2016, GO Biological Process 2018, GO Molecular Function 2018, and GO Cellular Component 2018 were used. Genes were ranked by -log_10_(p-value), and log_10_(OR). Benjamini-Hochberg correction was applied to p-values.

## Supplementary methods

### Immunofluorescence

Primary human monocyte derived macrophages were differentiated, as described above, on glass coverslips. Cells were infected as described. At 0, 2, 7, and 24 hours post infection cells were fixed for 20 min in 4% formaldehyde in PBS. After permeabilization with 0.2% Triton X-100 in PBS for 5 min at room temperature, cells were incubated with mouse monoclonal influenza A NP AA5H (BioRad) at 1:500. After 1 hour cells were washed three times with PBS and incubated with goat anti-mouse Alexa Fluor 488 at 1:1000 (ThermoFisher). After 1 hour cells were washed three times with PBS and incubated in DAPI (ThermoFisher) for ten minutes after which they were washed three times with PBS and mounted on slides using VECTASHIELD® Antifade Mounting Medium. Cells were viewed on a Leica fluorescence upright microscope and imaged using a Hamamatsu Orca-ER low light mono camera. Scale bars were added using ImageJ.

### Cell viability and Virus Titration

Cell viability was measured using Cell Titre Glo® at 0, 2, 7, and 24 hours post infection. Virus produced was titrated by plaque assay on MDCK cells. Virus titres in cell supernatants were determined by plaque titration using ten-fold serial dilutions of virus stocks. Confluent MDCK cells in 6 well plates were inoculated with cell supernatant for 1 hour in serum-free medium. An overlay (mixture of equal volume of DMEM and 2.4% Avicel (Sigma-Aldrich, UK) supplemented with 1 µg/ml TPCK-treated trypsin and 0.14% BSA fraction V) was then put onto the wells. After 48 hours, cells were fixed using 3.5% formaldehyde and stained with 0.1% crystal violet. Virus titres were calculated by plaque count*dilution factor/(volume of inoculum) and expressed as plaque forming units per millilitre of supernatant (pfu/ml).

### Identification of potential alternative splice variants

CAGE tags containing a leader sequence and an IAV promoter sequence followed by a sequence that did not align proximal to the IAV promoter sequence in the Udorn genome were extracted. These novel ‘promoter proximal’ sequences were hypothesised to be derived from putative 5’UTR sequences internal to a segment arising from mRNA from splice variants. These sequences were aligned throughout the Udorn genome using custom Python scripts. The abundance of each sequence was divided by the number of locations in the Udorn genome it could map to. The weighted abundances at each position were then summed and graphed. Segment 7 mRNA3 was used as a proof of principle.

### Determination of Coefficient of Variation Across 4 Donors

The expression profile of CTSS across all samples was extracted from the Fantom5 data in TPM (tags per million). The coefficient of variation for each gene was calculated for each of the 6 treatments across the 4 donors (SD:Mean * 100). Gene names were assigned to CTSS by Bedtools overlap with a window of +/- 5 bases using the hg19 annotation from Fantom5. Unnamed genes were removed from the final list. Results were separated by treatment.

## Supplementary Results

### Potential alternative splice variants

Spicing has been observed in segments 7 and 8 of IAV. In particular, in segment 7 the splice donor site for the mRNA3/M3 transcript is found at the end of the promoter sequence (Lamb, Lai and Choppin, 1981). Over 400,000 reads contained the IAV promoter sequence and a leader sequence, but did not originate from the genome sequence proximal to the promoter in any of the 8 segments. The leader and promoter sequences were removed and the sequences aligned throughout the Udorn genome. In order to quantify RNA expression at these loci, we summed the weighted abundances of reads originating at the same position. This revealed 6,902 putative capped IAV RNA sequences from the IAV genome, including the known splice variant of segment 7, the mRNA3 transcript (Figure S 3 D). The alignments observed (Table S 5) are likely to include previously unidentified splice variants. However, in a systematic search, no putative IAV splice variant RNA was preceded by a canonical major spliceosome acceptor site, apart from the mRNA3 transcript. It is possible these represent variants that are expressed in such low amounts they are not detectable by other means, for example northern blot or radioactive primer extension. It is of interest to determine if these putative mRNAs are true transcription products and if their transcription and translation contributes to viral pathogenesis.

## Supplementary Figures and Tables

**Figure S 1:**
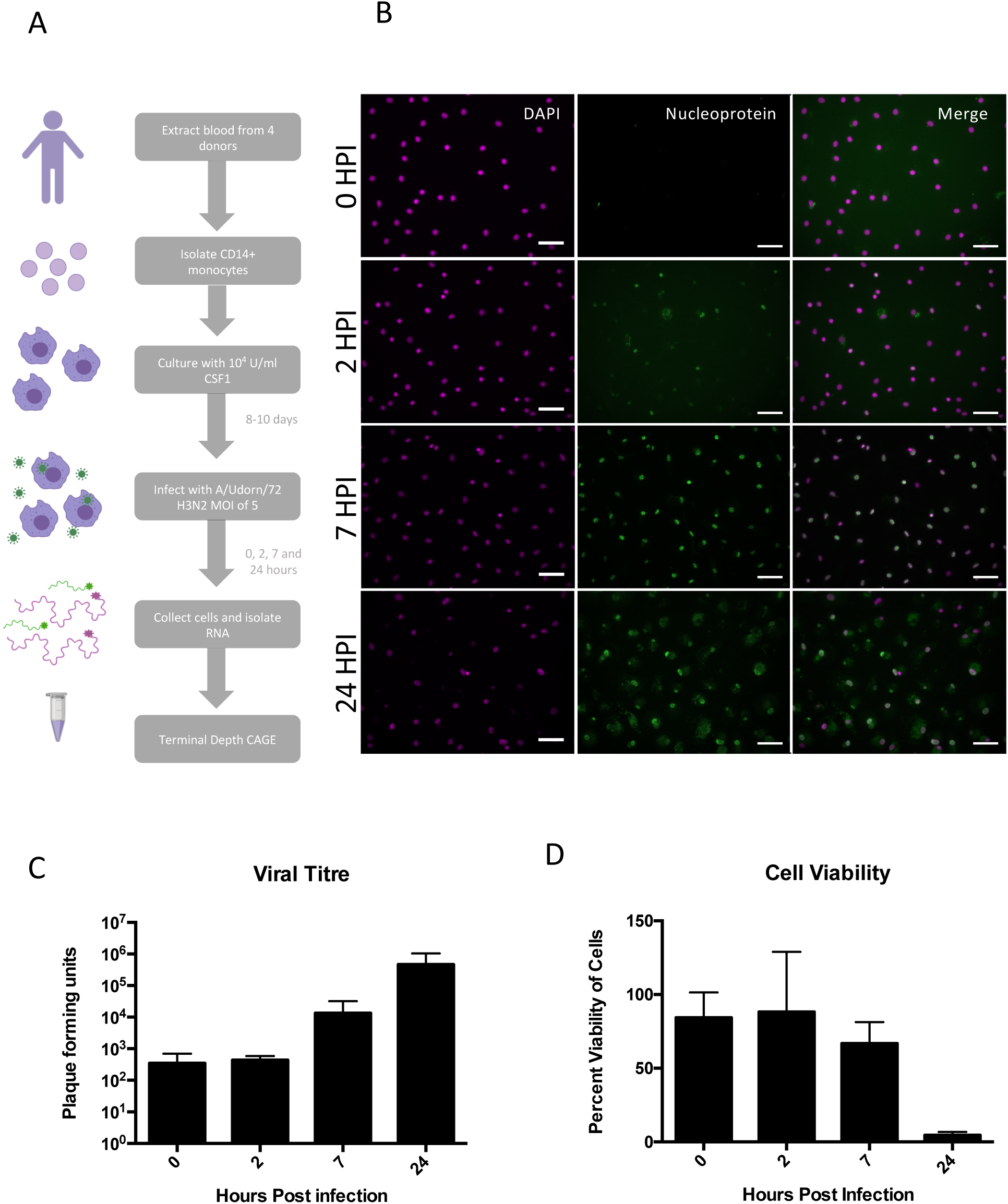
Characterisation of human monocyte derived macrophages productively infected with IAV. (A) Experimental outline. Blood was taken from 4 human donors, with appropriate ethical approval. CD14+ monocytes were extracted using magnetic beads and cultured in CSF1 for 8-10 days. MDMs were infected with A/Udorn/72 (H3N2) at a multiplicity of infection of 5. At 4 time points (0, 2, 7, and 24 hours after medium change) the cells were collected and RNA isolated. (B) Human MDMs were immunofluorescently stained for viral nucleoprotein to confirm infection at 0, 2, 7, and 24 hours post infection. Scale bars 10μm. (C) Viral titre was measured by plaque assay at 0, 2, 7, and 24 hours post infection (n = 3 independent experiments) and shown in pfu/ml supernatant. (D) Cell viability was measured using Cell Titre Glo® at 0, 2, 7, and 24 hours post infection (n = 3 independent experiments).

**Figure S 2:**
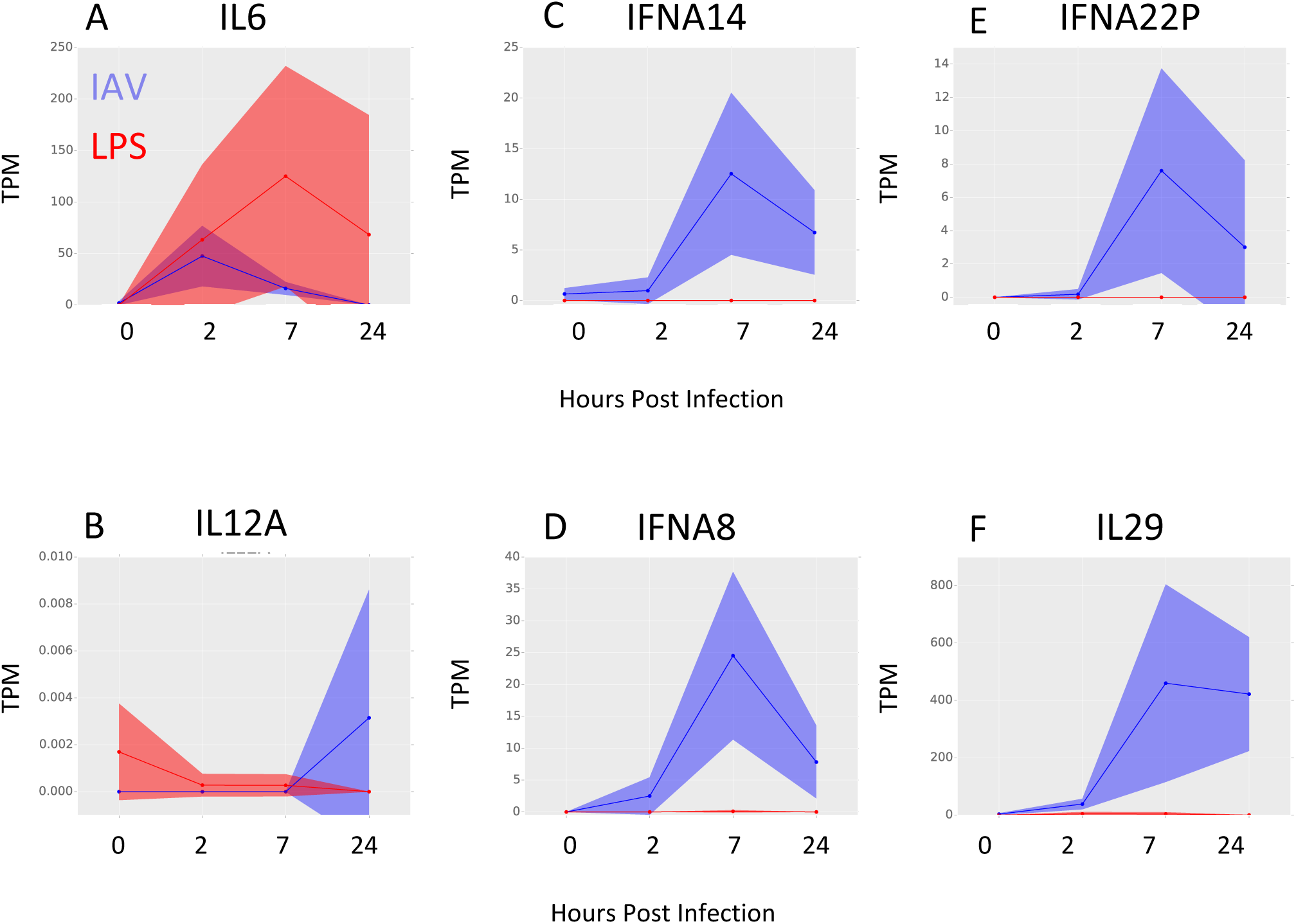
The transcriptional landscape between individual donors in response to IAV. (A-B) Comparison of the temporal response of transcripts between IAV- and LPS-treated MDMs. Relative expression of selected genes in LPS-treated (red) and IAV-infected (blue) human MDMs at 0, 2, 7, and 24 hours post treatment is shown in tags per million (TPM). Solid lines show the mean expression of all donors, filled-in area shows standard deviation between donors. (n = 3 for LPS, n = 4 for IAV). Filled-in area shows standard deviation between donors.

**Figure S 3:**
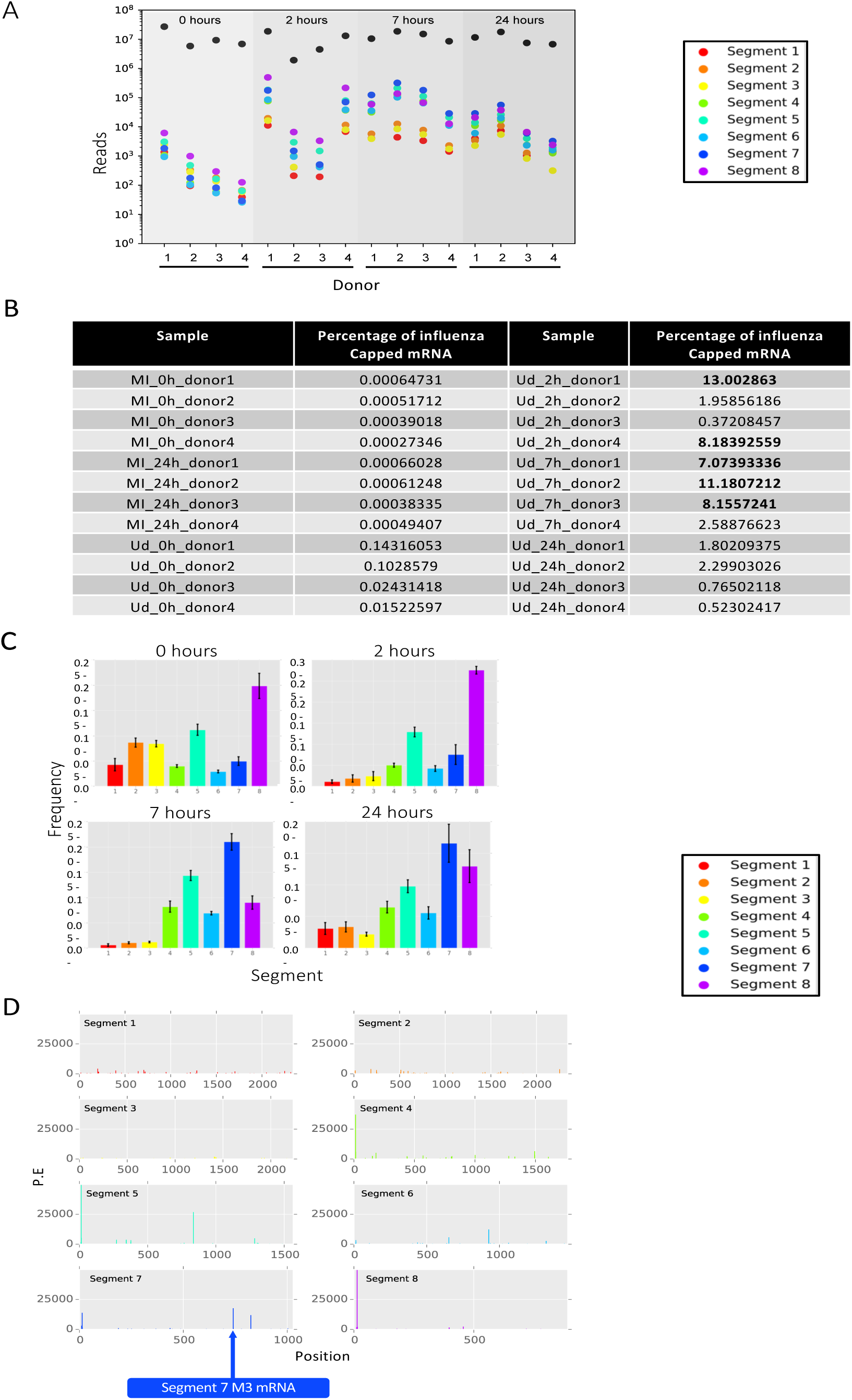
The amounts of IAV mRNA differed substantially between donors. (A) The raw number of IAV promoter-containing CAGE tags were counted, separated by segment sequence (see legend and below), shown alongside to those in the same sample that did not contain the viral promoter sequence (black). (B) Frequency, as percentage, of IAV promoter-containing CAGE tags in each sample. (C) The relative amount, compared to the total amount of viral mRNA, of mRNA from each viral segment was calculated for individual donors at each of the four timepoints. Height of the bar represents the mean frequency between donors. Error bars show standard deviation. (D) The positions of potential splice variant sequences aligned to the Udorn genome are shown as adjusted abundance. The known mRNA3 splice variant in segment 7 is shown (blue arrow). Time points and donors have been collated to increase signal.

**Figure S 4:**
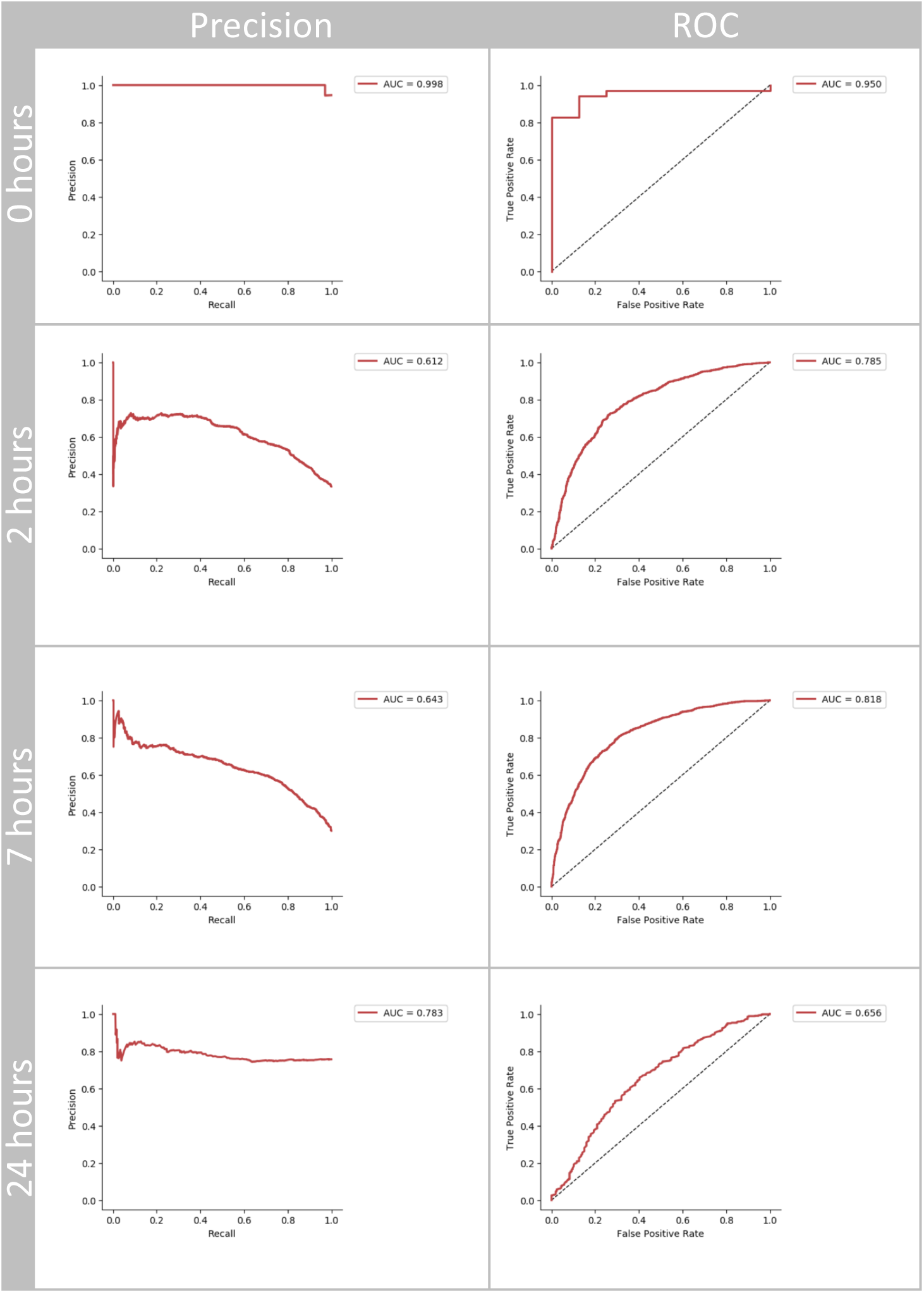
Precision Recall and Receiver Operator Curves for convolutional neural network analysis of 4 nucleotide length motifs in 10mer datasets. Models were trained and evaluated according to their receiver operating characteristic (ROC) area under the curve and Precision Recall area under the curve. The values for total area under the curve (AUC) are given. Each of the four time points was calculated independently.

**Figure S 5:**
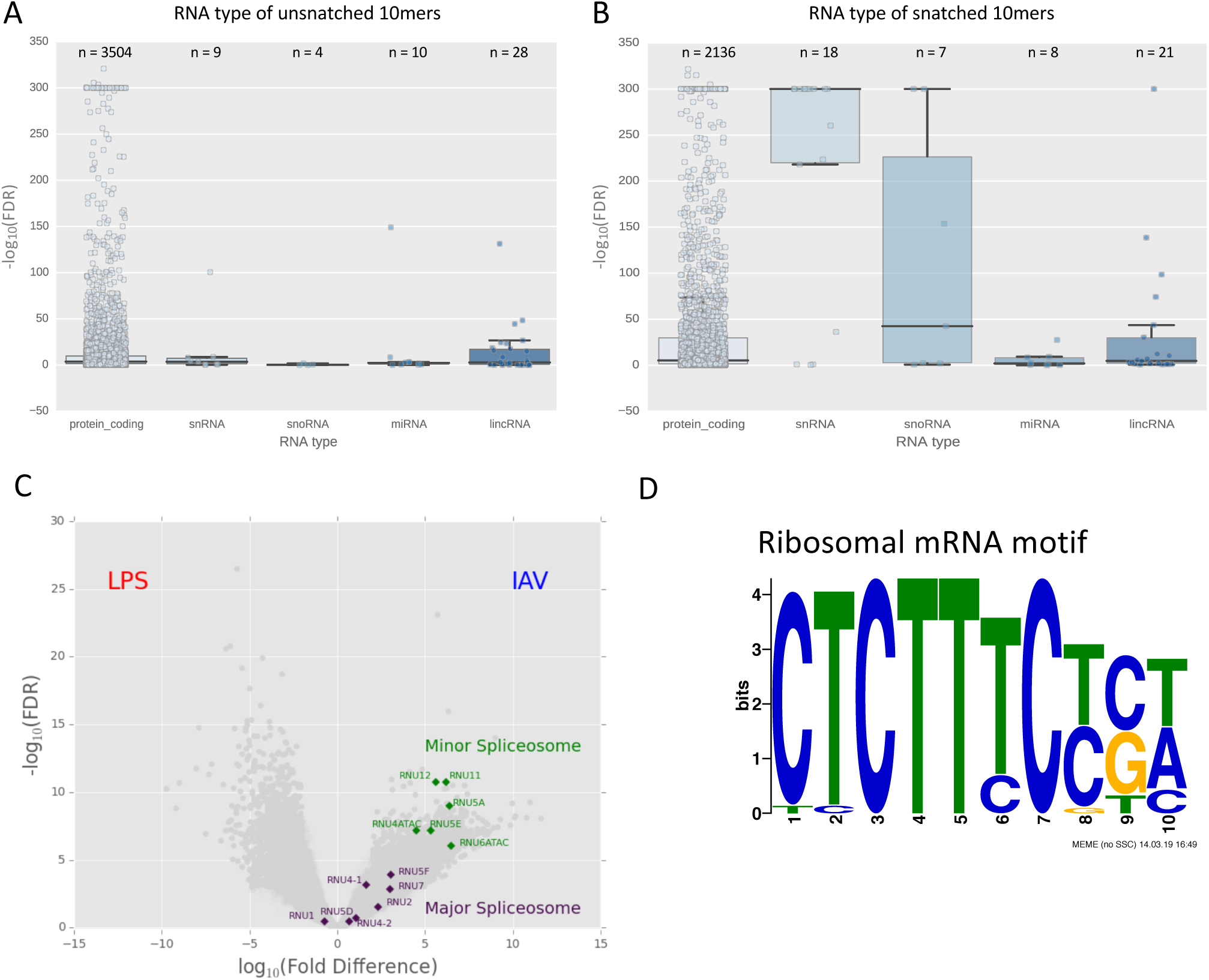
The IAV polymerase does snatch based on RNA type. (A, B) RNA type was assigned to 10mers based on transcript identity. Only 10mers with transcript identity were included. The significance of RNA type snatching was compared using ANOVA. RNA types were plotted against – log10(FDR) for 10mers of that type. The box denotes the interquartile range. The within the box represents the average and the whiskers represent standard deviation. The individual data-point for each 10mer is also plotted. The number of 10mers attributed to each RNA type is given as n above the box. (C) Differential gene expression analysis comparing expression of transcripts in LPS treated and IAV treated monocyte derived MDMs. Transcripts with a relative log fold change (log_2_FC) ≥ 3 and a -log_10_(FDR) ≥ 5 are shown in red (higher in LPS treated) and blue (higher in IAV infection). Components of the minor spliceosome are shown in green, while components of the major spliceosome are shown in purple. (D) MEME analysis of 10mers taken from human gene sequences corresponding to the first ten nucleotides of ribosomal protein mRNAs.

**Figure S 6:**
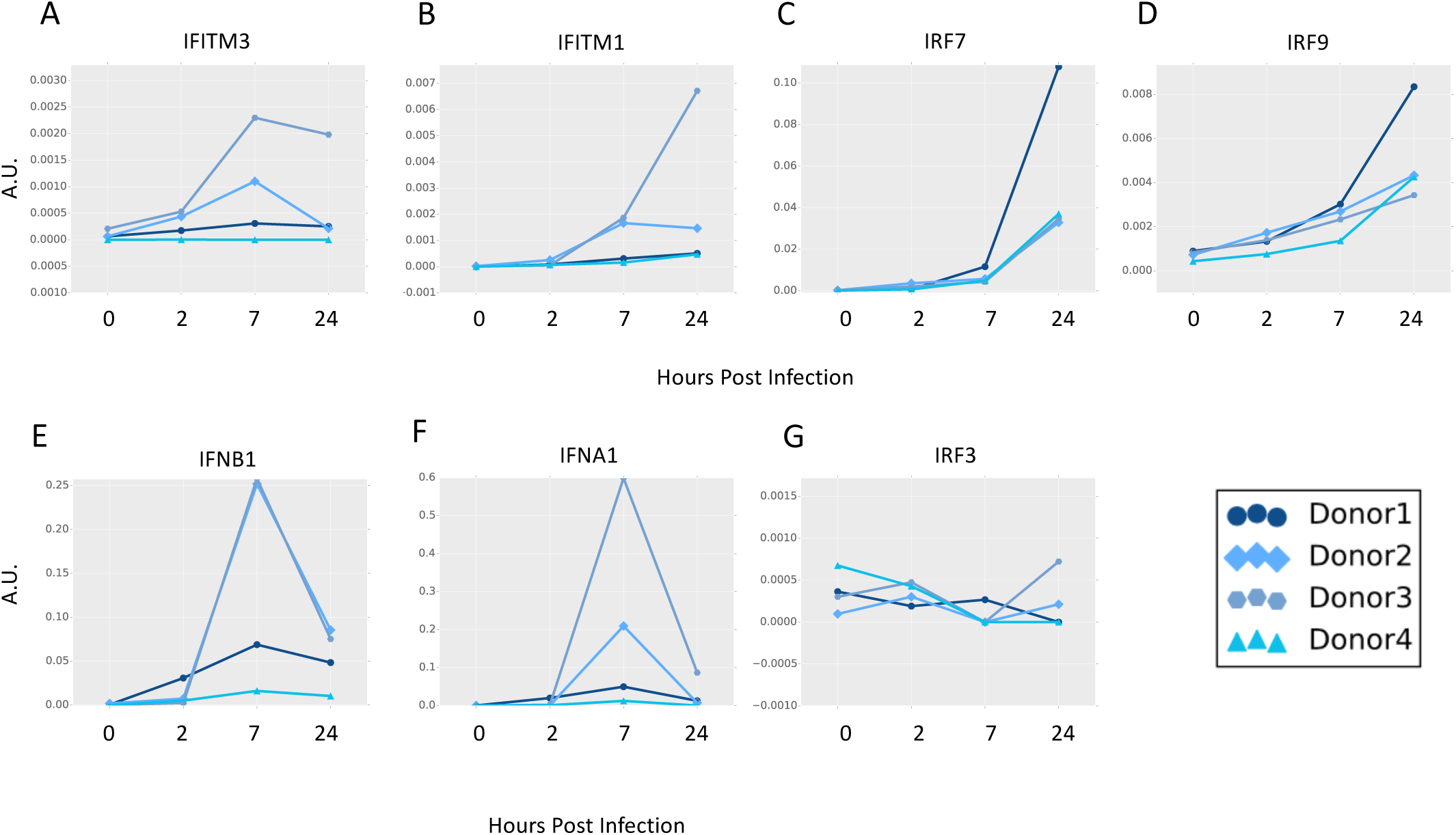
Variation in expression of interferon related genes between donors. Expression of the named genes at each of the four time points (0, 2, 7, and 24 hours post infection) was compared between donors (n=4). Expression is given in arbitrary units (A.U). Lines represent individual donors, as shown in legend.

## Supplementary Table Legends

**Table S 1: Clustering of transcripts expressed in IAV-infected MDMs.**

Analysis was restricted to the dominant promoters (p1) and used averaged data from the 4 donors. A Pearson correlation coefficient threshold of 0.94 and an MCL of 1.7 was used.

**Table S 2: GO term enrichment of the top 10 clusters.**

Analysis was performed using GATHER (Chang and Nevins, 2006).

**Table S 3: Enrichr analysis of the top 10 clusters.**

All results shown reach the threshold FDR < 0.5. Databases queried were Reactome 2016, KEGG 2016, and Wikipathways2016 (Kuleshov *et al*., 2016).

**Table S 4: Differential expression analysis of LPS- and IAV-treated MDMs.**

Analysis was performed using edgeR. All transcripts with a -log_2_(Fold Difference) >= 1 are shown.

**Table S 5: mRNA sequences for putative splice variants.**

**Table S 6:The most significantly and consistently snatched and unsnatched genes.**

All results shown reach the threshold FDR < 0.5.

**Table S 7: Fast preranked gene set enrichment analysis (FGSEA) of under- and over-represented 10mers.**

**Table S 8: Coefficient of variation for named genes at each time point.**

The coefficient of variation for named genes. Expression at CTSS were compared across donors (n=4) and annotated using publicly available Fantom5 annotation.

**Table S 9: Details of Udorn sequences used to assign identity to segments.**

Provided in Fasta format.

## Acknowledgements

The authors would like to thank Ross Hendry for advice on python code.

JKB gratefully acknowledges funding support from a Wellcome Trust Intermediate Clinical Fellowship (103258/Z/13/Z), a Wellcome-Beit Prize (103258/Z/13/A), and the UK Intensive Care Society. DAH, PD, and HW also acknowledge support from BBSRC Institute Strategic Programme Grants BB/J004324/1 and BB/P013740/1. KMS and DAH are supported by the Mater Foundation, Brisbane, Australia.

## Author Contributions

1. Conceptualization: SC JKB DAH PD NB HW KMS.

2. Data curation: JKB SC.

3. Formal analysis: SC JKB KMS DAH PD BW NP.

4. Funding acquisition: JKB DAH YH.

5. Investigation: SC JKB DAH PD KMS.

6. Methodology: SC JKB DAH.

7. Project administration: JKB DAH YH PC ARRF.

8. Resources: JKB DAH.

9. Supervision: JKB DAH YH PC ARRF.

10. Writing – original draft: SC JKB DAH KMS.

## Bibliography

Andersson, R. et al. (2014) ‘An atlas of active enhancers across human cell types and tissues’, Nature, 507(7493), pp. 455–461. doi: 10.1038/nature12787.

Bailey, T. L. and Elkan, C. (1994) ‘Fitting a mixture model by expectation maximization to discover motifs in biopolymers.’, Proceedings. International Conference on Intelligent Systems for Molecular Biology, 2(2), pp. 28–36. Available at: http://www.ncbi.nlm.nih.gov/pubmed/7584402.

Baillie, J. K. et al. (2016) ‘Shared activity patterns arising at genetic susceptibility loci reveal underlying genomic and cellular architecture of human disease.’, bioRxiv, pp. 1–24. doi: 10.1101/095349.

Baillie, J. K. et al. (2017) ‘Analysis of the human monocyte-derived macrophage transcriptome and response to lipopolysaccharide provides new insights into genetic aetiology of inflammatory bowel disease’, PLoS Genetics, 13(3), pp. 1–36. doi: 10.1371/journal.pgen.1006641.

Beaton, A. R. and Krug, R. M. (1981) ‘Selected host cell capped RNA fragments prime influenza viral RNA transcription in vivo’, Nucleic Acids Research, 9(17), pp. 4423–4436. doi: 10.1093/nar/9.17.4423.

Bercovich-Kinori, A. et al. (2016) ‘A systematic view on influenza induced host shutoff’, eLife, 5(AUGUST), pp. 1–20. doi: 10.7554/eLife.18311.

Budach, S. and Marsico, A. (2018) ‘Pysster: Classification of biological sequences by learning sequence and structure motifs with convolutional neural networks’, Bioinformatics, 34(17), pp. 3035–3037. doi: 10.1093/bioinformatics/bty222.

Canella, D. et al. (2010) ‘Defining the RNA polymerase III transcriptome: Genome-wide localization of the RNA polymerase III transcription machinery in human cells’, Genome Research, 20(6), pp. 710–721. doi: 10.1101/gr.101337.109.

Chang, J. T. and Nevins, J. R. (2006) ‘GATHER: A systems approach to interpreting genomic signatures’, Bioinformatics, 22(23), pp. 2926–2933. doi: 10.1093/bioinformatics/btl483.

Chen, E. Y. et al. (2013) ‘Enrichr: Interactive and collaborative HTML5 gene list enrichment analysis tool’, BMC Bioinformatics, 14. doi: 10.1186/1471-2105-14-128.

Cline, T. D., Beck, D. and Bianchini, E. (2017) ‘Influenza virus replication in macrophages: Balancing protection and pathogenesis’, Journal of General Virology, 98(10), pp. 2401–2412. doi: 10.1099/jgv.0.000922.

Everitt, A. R. et al. (2012) ‘IFITM3 restricts the morbidity and mortality associated with influenza.’, Nature, 484(7395), pp. 519–23. doi: 10.1038/nature10921.

Fairfax, B. P. et al. (2014) ‘Innate immune activity conditions the effect of regulatory variants upon monocyte gene expression’, Science, 343(6175). doi: 10.1126/science.1246949.

Forrest, A. R. R. et al. (2014) ‘A promoter-level mammalian expression atlas.’, Nature. Nature Publishing Group, 507(7493), pp. 462–70. doi: 10.1038/nature13182.

Freeman, T. C. et al. (2007) ‘Construction, visualisation, and clustering of transcription networks from microarray expression data’, PLoS Computational Biology, 3(10), pp. 2032–2042. doi: 10.1371/journal.pcbi.0030206.

Friesenhagen, J. et al. (2012) ‘Highly pathogenic avian influenza viruses inhibit effective immune responses of human blood-derived macrophages’, Journal of Leukocyte Biology, 92(1), pp. 11–20. doi: 10.1189/jlb.0911479.

Geerts-Dimitriadou, C., Goldbach, R. and Kormelink, R. (2011) ‘Preferential use of RNA leader sequences during influenza A transcription initiation in vivo’, Virology. Elsevier Inc., 409(1), pp. 27– 32. doi: 10.1016/j.virol.2010.09.006.

Gizzi, A. S. et al. (2018) ‘A naturally occurring antiviral ribonucleotide encoded by the human genome’, Nature. Springer US, 558(7711), pp. 610–614. doi: 10.1038/s41586-018-0238-4.

Gnirß, K. et al. (2015) ‘Tetherin Sensitivity of Influenza A Viruses Is Strain Specific: Role of Hemagglutinin and Neuraminidase’, Journal of Virology, 89(18), pp. 9178–9188. doi: 10.1128/JVI.00615-15.

Gu, W., Gallagher, Glen R, et al. (2015) ‘Influenza A virus preferentially snatches noncoding RNA caps.’, RNA (New York, N.Y.), 21(12), pp. 2067–75. doi: 10.1261/rna.054221.115.

Gu, W., Gallagher, Glen R., et al. (2015) ‘Influenza A virus preferentially snatches noncoding RNA caps’, Rna, 21(12), pp. 2067–2075. doi: 10.1261/rna.054221.115.

Haye, K. et al. (2009) ‘The NS1 Protein of a Human Influenza Virus Inhibits Type I Interferon Production and the Induction of Antiviral Responses in Primary Human Dendritic and Respiratory Epithelial Cells’, Journal of Virology, 83(13), pp. 6849–6862. doi: 10.1128/JVI.02323-08.

He, Y. et al. (2010) ‘Influenza A Virus Replication Induces Cell Cycle Arrest in G0/G1 Phase’, Journal of Virology, 84(24), pp. 12832–12840. doi: 10.1128/JVI.01216-10.

Hoeve, M. A. et al. (2012) ‘Influenza virus A infection of human monocyte and macrophage subpopulations reveals increased susceptibility associated with cell differentiation’, PLoS ONE, 7(1). doi: 10.1371/journal.pone.0029443.

Horby, P. et al. (2012) ‘The role of host genetics in susceptibility to influenza: A systematic review’, PLoS ONE, 7(3), pp. 1–9. doi: 10.1371/journal.pone.0033180. http://fantom.gsc.riken.jp/zenbu/ (no date) *Consortium, FANTOM 5*.

Hume, D. A. and Freeman, T. C. (2014) ‘Transcriptomic analysis of mononuclear phagocyte differentiation and activation.’, Immunological reviews, 262(1), pp. 74–84. doi: 10.1111/imr.12211.

Irvine, K. M. et al. (2009) ‘Colony-stimulating factor-1 (CSF-1) delivers a proatherogenic signal to human macrophages.’, Journal of leukocyte biology, 85(2), pp. 278–288. doi: 10.1189/jlb.0808497.

Jia, D. et al. (2010) ‘Influenza Virus Non-Structural Protein 1 (NS1) Disrupts Interferon Signaling’, PLoS ONE, 5(11), p. e13927. doi: 10.1371/journal.pone.0013927.

Kanamori-katayama, M. et al. (2011) ‘Unamplified cap analysis of gene expression on a single-molecule sequencer’, Cold Spring Harbor Genome, pp. 1150–1159. doi: 10.1101/gr.115469.110.enough.

Klinkhammer, J. et al. (2018) ‘IFN-λ prevents influenza virus spread from the upper airways to the lungs and limits virus transmission’, eLife, 7, p. e33354. doi: 10.7554/eLife.33354.

Koppstein, D., Ashour, J. and Bartel, D. P. (2015) ‘Sequencing the cap-snatching repertoire of H1N1 influenza provides insight into the mechanism of viral transcription initiation’, Nucleic Acids Research, 43(10), pp. 1–13. doi: 10.1093/nar/gkv333.

Koppstein, David, Ashour, J. and Bartel, D. P. (2015) ‘Sequencing the cap-snatching repertoire of H1N1 influenza provides insight into the mechanism of viral transcription initiation’, Nucleic Acids Research, 43(10), pp. 5052–5064. doi: 10.1093/nar/gkv333.

Kuleshov, M. V. et al. (2016) ‘Enrichr: a comprehensive gene set enrichment analysis web server 2016 update’, Nucleic acids research, 44(W1), pp. W90–W97. doi: 10.1093/nar/gkw377.

Lamb, R. A., Lai, C. J. and Choppin, P. W. (1981) ‘Sequences of mRNAs derived from genome RNA segment 7 of influenza virus: colinear and interrupted mRNAs code for overlapping proteins.’, Proceedings of the National Academy of Sciences of the United States of America, 78(7), pp. 4170– 4. doi: 10.1073/pnas.78.7.4170.

Lee, M. N. et al. (2014) ‘Common Genetic Variants Modulate Pathogen-Sensing Responses in Human Dendritic Cells’, Science, 343(6175), pp. 1246980–1246980. doi: 10.1126/science.1246980.

Lee, S. M. Y. et al. (2009) ‘Systems-level comparison of host-responses elicited by avian H5N1 and seasonal H1N1 influenza viruses in primary human macrophages’, PLoS ONE, 4(12). doi: 10.1371/journal.pone.0008072.

McCauley, J. W. and Mahy, B. W. (1983) ‘Structure and function of the influenza virus genome.’, The Biochemical journal, 211(2), pp. 281–94. doi: 10.1042/bj2110281.

Monteerarat, Y. et al. (2010) ‘Induction of TNF-α in human macrophages by avian and human influenza viruses’, Archives of Virology, 155(8), pp. 1273–1279. doi: 10.1007/s00705-010-0716-y.

Morita, E. et al. (2007) ‘Identification of Human MVB12 Proteins as ESCRT-I Subunits that Function in HIV Budding’, Cell Host and Microbe, 2(1), pp. 41–53. doi: 10.1016/j.chom.2007.06.003.

Morris, D. E., Cleary, D. W. and Clarke, S. C. (2017) ‘Secondary bacterial infections associated with influenza pandemics’, Frontiers in Microbiology, 8(JUN), pp. 1–17. doi: 10.3389/fmicb.2017.01041.

Nicol, M. Q. and Dutia, B. M. (2014) ‘The role of macrophages in influenza A virus infection’, Future Virology, 9(9), pp. 847–862. doi: 10.2217/fvl.14.65.

Nyman, T. a et al. (2000) ‘Proteome analysis reveals ubiquitin-conjugating enzymes to be a new family of interferon-alpha-regulated genes.’, European journal of biochemistry / FEBS, 267(13), pp. 4011–9. Available at: http://www.ncbi.nlm.nih.gov/pubmed/10866800.

Ohman, T. et al. (2009) ‘Actin and RIG-I/MAVS Signaling Components Translocate to Mitochondria upon Influenza A Virus Infection of Human Primary Macrophages’, The Journal of Immunology, 182(9), pp. 5682–5692. doi: 10.4049/jimmunol.0803093.

Perez-Cidoncha, M. et al. (2014) ‘An Unbiased Genetic Screen Reveals the Polygenic Nature of the Influenza Virus Anti-Interferon Response’, Journal of Virology, 88(9), pp. 4632–4646. doi: 10.1128/JVI.00014-14.

Perrone, L. A. et al. (2008) ‘H5N1 and 1918 pandemic influenza virus infection results in early and excessive infiltration of macrophages and neutrophils in the lungs of mice’, PLoS Pathogens, 4(8). doi: 10.1371/journal.ppat.1000115.

Plotch, S. J. et al. (1981) ‘A unique cap(m7GpppXm)-dependent influenza virion endonuclease cleaves capped RNAs to generate the primers that initiate viral RNA transcription’, Cell, 23(3), pp. 847–858. doi: 10.1016/0092-8674(81)90449-9.

Ramilo, O. et al. (2018) ‘Gene expression patterns in blood leukocytes discriminate patients with acute infections’, 109(5), pp. 1–2. doi: 10.1182/blood-2006-02-002477.The.

Rao, P., Yuan, W. and Krug, R. M. (2003) ‘Crucial role of CA cleavage sites in the cap-snatching mechanism for initiating viral mRNA synthesis’, EMBO Journal, 22(5), pp. 1188–1198. doi: 10.1093/emboj/cdg109.

van Riel, D. et al. (2011) ‘Highly pathogenic avian influenza virus H5N1 infects alveolar macrophages without virus production or excessive TNF-alpha induction’, PLoS Pathogens, 7(6), pp. 4–11. doi: 10.1371/journal.ppat.1002099.

Van Riel, D. et al. (2007) ‘Human and avian influenza viruses target different cells in the lower respiratory tract of humans and other mammals’, American Journal of Pathology, 171(4), pp. 1215–1223. doi: 10.2353/ajpath.2007.070248.

Robinson, M. D., McCarthy, D. J. and Smyth, G. K. (2009) ‘edgeR: A Bioconductor package for differential expression analysis of digital gene expression data’, Bioinformatics, 26(1), pp. 139–140. doi: 10.1093/bioinformatics/btp616.

Sergushichev, A. (2016) ‘An algorithm for fast preranked gene set enrichment analysis using cumulative statistic calculation’, bioRxiv, p. 60012. doi: 10.1101/060012.

Short, K. R. et al. (2017) ‘Proinflammatory Cytokine Responses in Extra-Respiratory Tissues during Severe Influenza’, Journal of Infectious Diseases, 216(7), pp. 829–833. doi: 10.1093/infdis/jix281.

Sikora, D. et al. (2014) ‘Deep sequencing reveals the eight facets of the influenza A/HongKong/1/1968 (H3N2) virus cap-snatching process.’, Scientific reports. doi: 10.1038/srep06181.

Sikora, D. et al. (2017) ‘Influenza A virus cap-snatches host RNAs based on their abundance early after infection’, Virology. Elsevier Inc., 509(June), pp. 167–177. doi: 10.1016/j.virol.2017.06.020.

Singh, R. and Reddy, R. (2006) ‘Gamma-monomethyl phosphate: a cap structure in spliceosomal U6 small nuclear RNA.’, Proceedings of the National Academy of Sciences, 86(21), pp. 8280–8283. doi: 10.1073/pnas.86.21.8280.

Söderholm, S. et al. (2016) ‘Phosphoproteomics to Characterize Host Response During Influenza A Virus Infection of Human Macrophages’, Molecular & Cellular Proteomics, 15(10), pp. 3203–3219. doi: 10.1074/mcp.M116.057984.

Stasakova, J. et al. (2005) ‘Influenza A mutant viruses with altered NS1 protein function provoke caspase-1 activation in primary human macrophages, resulting in fast apoptosis and release of high levels of interleukins 1β and 18’, Journal of General Virology, 86(1), pp. 185–195. doi: 10.1099/vir.0.80422-0.

Takahashi, H. et al. (2012) ‘5’ end-centered expression profiling using cap-analysis gene expression and next-generation sequencing. TL - 7’, Nature protocols. Nature Publishing Group, 7 VN-re(3), pp. 542–561. doi: 10.1038/nprot.2012.005.

Teijaro, J. R. (2014) ‘The Role of Cytokine Responses During Influenza Virus Pathogenesis and Potential Therapeutic Options’, in Influenza Pathogenesis and Control - Volume II, pp. 3–22.

Thakar, J. et al. (2013) ‘Overcoming NS1-Mediated Immune Antagonism Involves Both Interferon-Dependent and Independent Mechanisms’, Journal of Interferon & Cytokine Research, 33(11), pp. 700–708. doi: 10.1089/jir.2012.0113.

De Vlugt, C., Sikora, D. and Pelchat, M. (2018) ‘Insight into Influenza: A Virus Cap-Snatching’, Viruses, 10(11), p. 641. doi: 10.3390/v10110641.

Wang, J. et al. (2012) ‘Innate immune response of human alveolar macrophages during influenza a infection’, PLoS ONE, 7(3). doi: 10.1371/journal.pone.0029879.

Wang, W. and Krug, R. M. (1998) ‘U6atac snRNA, the highly divergent counterpart of U6 snRNA, is the specific target that mediates inhibition of AT-AC splicing by the influenza virus NS1 protein.’, RNA (New York, N.Y.), 4(1), pp. 55–64. Available at: http://www.pubmedcentral.nih.gov/articlerender.fcgi?artid=1369596&tool=pmcentrez&rendertype=abstract.

Wang, X., Hinson, E. R. and Cresswell, P. (2007) ‘The Interferon-Inducible Protein Viperin Inhibits Influenza Virus Release by Perturbing Lipid Rafts’, Cell Host and Microbe, 2(2), pp. 96–105. doi: 10.1016/j.chom.2007.06.009.

Wei, J. et al. (2019) ‘Ribosomal Proteins Regulate MHC Class I Peptide Generation for Immunosurveillance’, Molecular Cell, pp. 1162–1173. doi: 10.1016/j.molcel.2018.12.020.

WHO (no date) http://www.who.int/mediacentre/factsheets/fs211/en/.

Younis, I. et al. (2013) ‘Minor introns are embedded molecular switches regulated by highly unstable U6atac snRNA’, eLife, 2013(2), pp. 1–14. doi: 10.7554/eLife.00780.

